# B Cells Promote T Cell Immunosenescence and Mammalian Aging Parameters

**DOI:** 10.1101/2023.09.12.556363

**Authors:** Saad Khan, Mainak Chakraborty, Fei Wu, Nan Chen, Tao Wang, Yi Tao Chan, Azin Sayad, Juan Diego Sánchez Vásquez, Max Kotlyar, Khiem Nguyen, Yingxiang Huang, Faisal J. Alibhai, Minna Woo, Ren-Ke Li, Mansoor Husain, Igor Jurisica, Adam J. Gehring, Pamela S. Ohashi, David Furman, Sue Tsai, Shawn Winer, Daniel A. Winer

## Abstract

A dysregulated adaptive immune system is a key feature of aging, and is associated with age-related chronic diseases and mortality. Most notably, aging is linked to a loss in the diversity of the T cell repertoire and expansion of activated inflammatory age-related T cell subsets, though the main drivers of these processes are largely unknown. Here, we find that T cell aging is directly influenced by B cells. Using multiple models of B cell manipulation and single-cell omics, we find B cells to be a major cell type that is largely responsible for the age-related reduction of naive T cells, their associated differentiation towards pathogenic immunosenescent T cell subsets, and for the clonal restriction of their T cell receptor (TCR). Accordingly, we find that these pathogenic shifts can be therapeutically targeted via CD20 monoclonal antibody treatment. Mechanistically, we uncover a new role for insulin receptor signaling in influencing age-related B cell pathogenicity that in turn induces T cell dysfunction and a decline in healthspan parameters. These results establish B cells as a pivotal force contributing to age-associated adaptive immune dysfunction and healthspan outcomes, and suggest new modalities to manage aging and related multi-morbidity.

**One Sentence Summary:** Insulin receptor signaling facilitates the induction of age associated B cell inflammatory changes, which drive phenotypic aging of the T cell compartment and adverse outcomes to mammalian healthspan parameters.

## Introduction

Aging is associated with a gradual decline in tissue and organ function, often resulting in the development of chronic complications and diminished survivability (*1, 2*). The unprecedented growth of the global aged population (>65 years) is of great concern, as the elderly display the highest death rate linked with non-communicable diseases, such as cardiovascular disease, type 2 diabetes, and Alzheimer’s disease (*3*). Consequently, there is great interest in identifying potential therapeutics that can prevent age-related decline in human health.

Mechanisms that drive age-related dysfunction occur in tandem, influence one another, and overlap significantly in their effect on disease manifestation; these mechanisms are collectively referred to as the pillars and hallmarks of aging (*2, 4*). One of these pillars, is the development of a state of chronic low-grade inflammation, termed “inflammaging”, that progressively increases and is directly linked with the onset of age-related complications and decreased survival of the elderly (*5*). Cells of the immune system are central to the instigation and progression of inflammaging, and age-related changes to the immune system are collectively termed “immunosenescence” (*6*). Indeed, immunosenescence is directly responsible for age-related increases in basal levels of inflammation, decreased vaccine efficacy, and increased susceptibility to pathogenic agents, such as the SARS-CoV2 coronavirus (*6, 7*). As such, there is broad interest in identifying shifts within the immune system that occur with age and the mechanisms that drive them.

The adaptive arm of the immune system, comprising of B cells and T cells, is reflective of the immunosenescence process and is the primary cause of an age-related decline in the generation of immunological memory against infectious agents. Most notable are alterations to the T cell compartment, which display a loss in their antigen-inexperienced naive pool and an expansion of memory T cells, the latter of which possess an elevated inflammatory tone capable of driving inflammaging (*8, 9*). Furthermore, the imbalance in the ratio of the naive/memory T cells is associated with a loss in T cell receptor (TCR) diversity that potentially compromises responses to foreign agents (*8, 9*). The current thinking is that thymic involution and hematopoietic stem cell insufficiency are partly responsible for the loss in naive T cells with age, but these points are controversial as homeostatic proliferation and maintenance may be the main force controlling the T cell repertoire during aging (*9, 10*). Thus, defects in biological systems regulating peripheral maintenance of T cells are likely responsible, yet what these mechanistic axes are remain largely understudied. Moreover, it is unclear if changes in the T cell repertoire with age are due to defects in the T cells themselves, or if these changes are a consequence of crosstalk with other cell types.

Aging is associated with a decline in early B cell progenitors, and a similar shift away from a naive dominant compartment towards one that displays greater features associated with activation, which includes increased production of pro-inflammatory cytokines, memory-associated phenotypes, and terminal differentiation towards antibody secreting cells (ASCs) (*11–13*). Indeed, epidemiological evidence suggests that aging is linked with increased titers of non-organ specific autoantibodies and compromised humoral immunity upon vaccination (*12*). One of the most noticeable alterations to the aging B cell compartment is the formation of a unique B cell subset termed age-associated B cells (ABCs), that in mice lack expression of CD21 and CD23, display expression of CD11c and the transcription factor T-bet (*14*). ABCs reportedly express higher levels of pro-inflammatory cytokines, the antibody IgG2a/c, and antigen-presenting molecules, and as such, are likely integral to the development of inflammaging and immunosenescence (*14*).

Collectively, B cells are central to dictating immune responses to infection during mammalian health and have been implicated in contributing to metabolic disease and autoimmunity (*15–17*). Here, we show that B cells are critical in dictating overall adaptive immune dysfunction with age and are a major cell type that drives age associated changes to T cell composition, exhaustion, inflammatory potential, as well as the altered TCR repertoire in aged mice. B cells undergo distinct phenotypic changes with age, in line with an immunosenescent phenotype. As part of this phenotype, we find that B cells can engage in crosstalk with T cells to influence their fate during aging, and further affect systemic parameters related to mammalian healthspan. Mechanistically, B cells dictate T cell parameters at the transcriptional level which in turn is responsible for alterations to TCR diversity and T cell clonality with age. We demonstrate that B cells can be targeted therapeutically to curb these consequences, and thus identify the potential for using B cell targeted therapies to prevent age-related complications. Finally, we uncover a novel mechanistic role for the insulin receptor (InsR) in mediating its pro-aging effects on mice by acting on B cells to promote age associated B cell phenotypes that drive T cell immunosenescence and reduced healthspan. Overall, these findings link an evolutionary conserved endocrine signaling axis with dysfunctional adaptive immunity and declining health parameters with age.

## Results

### Aging alters T and B cell compartments in mice and humans

We first explored alterations to the composition of the splenic adaptive immune compartment in 22–30-month-old (aged) C57BL6/J wild type (WT) mice (gating strategy: Fig. S1). In comparison to 3-6-month-old (young) WT mice, aged mice displayed a slight decrease in the relative abundance of CD19^+^ B cells, yet their overall cellularity and composition of B220^lo/−^ and B220^hi^ subsets remained unchanged (Fig. S2A-B). CD19^+^ B220^hi^ (B2) cells are the majority B cell fraction that are observed in mice and humans and are essential for humoral immunity, antigen presentation to T cells, and generation of immunological memory (*18–20*). Within the B2 fraction, we noticed an age-associated decline in the abundance of follicular B cells (FOBs) and a concurrent expansion of a CD21^−^ CD23^−^ fraction, previously described as ABCs (*19*) (Fig. 1A). Aging has also been associated with an expansion of CD11c^+^ T-bet^+^ B cells (*14*); indeed, we observed that this phenotype was exclusive to the B2 compartment and accordingly, CD11c^+^ T-bet^+^ B2 cells displayed increased cellularity within the spleen (Fig. 1B). Additionally, within the B2 cell subset, we observed an increase in IgM^−^ IgD^−^ cells undergone class switch recombination (CSR), and memory-like CD80^+^ PDL2^+^ cells (Fig. 1C), implying potential antigen encounter by B2 cells during the aging process.

**Fig. 1:**
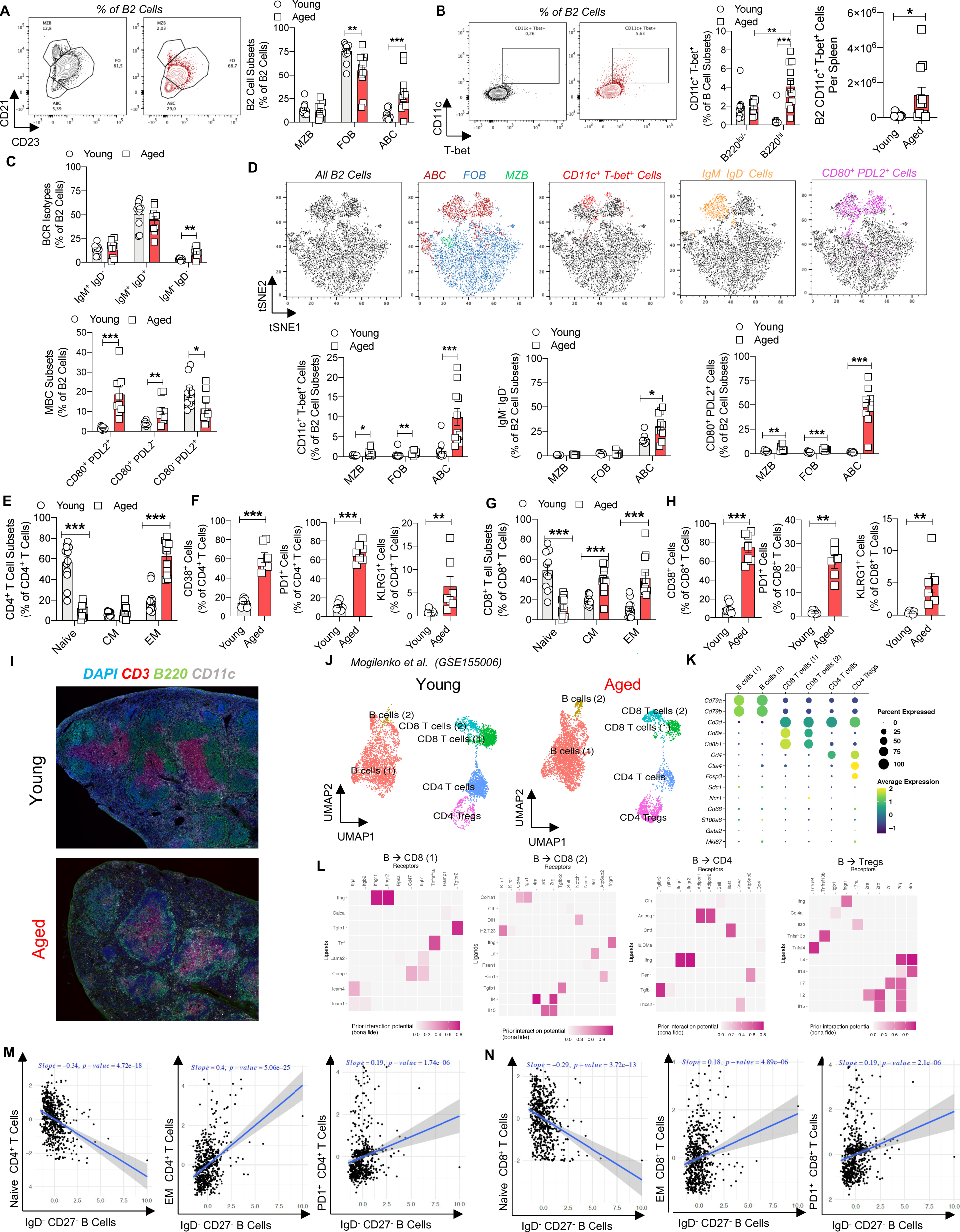
Age-related changes to B and T cells is associated with increased cellular crosstalk. (A-H): Flow cytometric assessment of splenic B and T cells in young and aged WT mice (n= 9-12 per group). (A) Representative plots (left) and relative abundance (right) of splenic B2 cell subsets. (B) Representative plots of CD11c^+^ T-bet^+^ B2 cells (left), relative abundance of CD11c^+^ T-bet^+^ B cells (middle) and cell number of CD11c^+^ T-bet^+^ B2 cells. (C)Abundance of B2 B cell receptor isotypes (top) and memory B cell subsets (bottom). (D) Representative tSNE plots of B2 cell subsets (top) and abundance of CD11c^+^ T-bet^+^ cells (bottom left), IgM^−^ IgD^−^ cells (bottom middle) and CD80^+^ PDL2^+^ cells (bottom right). (E) Naive, effector memory, and central memory abundance in CD4^+^ T cells. (F) CD38^+^ cells (left), PD1^+^ cells (middle) and KLRG1^+^ cells (right) abundance in CD4^+^ T cells. (G) Naive, effector memory, and central memory abundance in CD8^+^ T cells. (H) CD38^+^ cells (left), PD1^+^ cells (middle) and KLRG1^+^ cells (right) abundance in CD8^+^ T cells (I) Representative immunofluorescent staining of CD3, B220 and CD11c in young (top) and aged (bottom) spleen. (J-L): NicheNet analysis on B cell sender populations driving age-related T cell changes (Mogilenko *et al.* GSE155006). (J) UMAP of young (left) and aged (right) re-analyzed splenic B and T cell cluster. (K) Gene expression profile of individual clusters from aggregated young and aged splenic samples. (L) NicheNet predictive analysis highlighting B cell ligands interacting with various T cell cluster receptors driving age-related changes to T cells. (M-N) Correlation of IgD^−^ CD27^−^ B cells with T cell subsets in peripheral blood of 1000 immunomes cohort (n=609). (M) Correlation of naive (left), effector memory (middle) and PD1^+^ (right) CD4^+^ T cells against IgD^−^ CD27^−^ B cells. (N) Correlation of naive (left), effector memory (middle) and PD1^+^ (right) CD8^+^ T cells against IgD^−^ CD27^−^ B cells. Data are means ± SEM. * denotes *p* < 0.05, ** denotes *p* < 0.01, and *** denotes *p* < 0.001.

We next sought to understand which B2 cell subset is responsible for these age-related differences. Remarkably, the expansion of effector CD11c^+^ T-bet^+^ cells, class-switched IgM^−^ IgD^−^ cells, and memory-like CD80^+^ PDL2^+^ cells, was more pronounced within the ABC compartment, compared to FOB and MZB cells (Fig. 1D). Interestingly, we also observed a decrease in IgM^+^ IgD^−^ cells and an increase in CD80^+^ PDL2^−^ cells exclusively within the ABC compartment (Fig. S2C-D). An increased memory like phenotype suggests ABCs may in themselves represent a pool of antigen-experienced B cells that become increasingly activated and memory-like with age. Separately, within the splenic B220^lo/−^ compartment we observed no difference in the abundance of CD5^+^ and CD5^−^ subsets, or BCR isotype expression patterns; we did however note increases in memory cell subsets (Fig. S2E)

Compared to young controls, aged mice displayed no difference in the frequency or cellularity of total splenic T cells, although, aging did result in reduced abundance of CD8^+^ T cells (Fig. S2F-G). Within both CD4^+^ and CD8^+^ compartments we observed a marked decrease in the abundance of naive T cells and an expansion of effector memory (EM) T cells (Fig. 1E, G). CD8+ T cells also displayed an additional increase in the abundance of central memory (CM) T cells (Fig. 1G). Additional profiling unveiled an expansion of PD1^+^, CD38^+^, and KLRG1^+^ T cells with age (Fig. 1F, H), which have traditionally been linked with exhaustion, activation, and senescence, respectively.

Immunofluorescent assessment of the spleen revealed that aging was associated with a loss in clear segregation of splenic B cell and T cell zones (Fig. 1I), in line with previous reports regarding disruptions to microanatomy of the spleen with age (*21*). As a result, B cells and T cell interactions appeared to take place outside of traditional B-T cell borders, which might result in greater crosstalk amongst the two cells with age. To understand whether B cells could directly influence the age-related phenotype of T cells within the spleen, we assessed a previously published splenic single cell RNA sequencing dataset (*22*). After reprocessing, quality control, and normalization we sub-clustered B cells, CD8^+^ T cells, and CD4^+^ T cells from a heterogeneous splenic compartment (Fig. S2H-I). Upon sub-clustering, we identified two B cell clusters, two CD8^+^ T cell clusters, a CD4^+^ T cell cluster and a regulatory T cell cluster (Fig. 1J-K and Fig. S2J-K). Of note, B cell cluster 2 appeared to expand with age, lacked expression of CD21 (*Cr2)* and CD23 (*Fcer2a)*, and displayed selective expression of CD11c (*Itgax*) and Tbet (*Tbx21*) (Fig. S2L), akin to the previously identified ABCs. To examine intercellular crosstalk, we next applied NicheNet analyses (*23*) and designated both B cell clusters as sender populations and all T cell clusters as receivers, with the aged spleen set as the affected condition and the young spleen set as the control. B cells were thereby predicted to potentially age the T cell compartment via production of secreted factors (e.g., *Ifng, Tnf, Tgfb1*) and expression of surface proteins (e.g., *Icam1, H2T23, H2.DMa*), which also differed in expression among the 2 B cell clusters, implying potentially non-redundant roles in influencing T cell aging (Fig. 1L and Fig. S2M-P).

We subsequently assessed alterations to B cells and T cells in the blood (Fig. S3A-H), liver (Fig. S3I-P), and lungs (Fig. S3Q-X) of young and aged WT mice by flow cytometry. Our findings were similar to the spleen, where aging resulted in B cells adopting an age-associated phenotype that was concurrent with an expansion of pathogenic T cell subsets and a loss in naive T cells (Fig. S3A-X). Pathway analysis of a previously published dataset containing young and aged splenic, hepatic, pulmonary, and peritoneal B cells revealed that aging upregulated pathways related to increased B cell activation, antigen presentation, cytokine production and T cell polarization capabilities (Fig. S3Y). We further observed that in some instances, B cells and T cells collaboratively formed ectopic tertiary lymphoid structures (TLS) within aged livers and the lungs, which could represent potential hubs for interactions outside of lymphoid organs (Fig. S3Z).

We next determined whether our findings could be applicable to human aging. We first re-analyzed the Tabula Sapiens dataset (*24*), to assess crosstalk amongst B cells and T cells in the aged human spleen. Like our mouse NicheNet analyses, aged B cells were observed to be capable of interacting with aged T cells via their production of secreted and surface proteins (e.g., *IL-15, TGFB1, TNF, CD40, HLA.DMA, HLA.A, ICAM1*) (Fig. S4A-E), confirming this communication axis is conserved in humans. In collaboration with the 1000 immunomes project (*25*), we sought to specifically assess the relevance of a human ABC compartment in the peripheral blood of individuals across the human lifespan (Fig. S4F). In humans, ABCs are demarcated by their lack of expression of IgD and CD27, in addition to displaying similar expression patterns of CD11c and T-bet, CSR, and memory phenotypes as mice (*14*). First, we found that the abundance of IgD^−^ CD27^−^ B cells was positively correlated with the inflammatory age (iAge) of humans (Fig. S4G), a metric for chronic age-related inflammation in humans linked with immunosenescence, frailty, and multimorbidity (*25*). In corroboration with our findings in mice, the abundance of human ABC-like cells correlated with an increase in effector memory, central memory, and PD1+ T cells, and a decrease in naive T cells within the blood of people at any age across the lifespan, within the 1000 immunomes cohort (Fig. 1M-N).

Our findings here are two-fold. First, alterations to the adaptive immune compartment are systemic, and observable in mice and humans; henceforth, they could serve as potential biomarkers of mammalian aging. Secondly, age-associated shifts in B cells occurs in congruence with alterations to T cells, and raises the idea of B cell-T cell crosstalk as a critical modulator of the T cell inflammaging process.

### B cell deficiency curtails T cell Inflammaging and age-associated multi-morbidity

B cells are key stimulatory cells that can drive T cell activation during infection, autoimmunity, and metabolic disease (*17, 26*). Based on our results from the previous experiments in young and aged WT mice, we rationalized that an absence of B cells might limit the expansion of age-associated pathogenic T cell subsets. Thus, we aged B cell deficient (μMT) mice, which lack a membrane-bound IgM and consequently display a B cell developmental arrest within the bone marrow at the pre-B cell stage, to approximately 24 months (*27*). As expected, spleens of aged μMT mice displayed reductions in the cellularity and relative abundance of total B cells (Fig. S5A). As a result, the relative abundance of splenic T cells increased in μMT mice, despite an overall similar decrease in cellularity, compared to WT controls (Fig. S5B). Additionally, within the T cell compartment, we noted an expansion in the abundance of CD8^+^ T cells and a reduction in the relative abundance of CD4^+^ T cells (Fig. S5C).

Remarkably, aged μMT mice displayed a splenic T cell compartment that was phenotypically resistant to age-related changes. Comparted to WT mice, μMT mice possessed increased abundances of naive T cells, along with marked reductions in the abundance of EM T cells and PD1^+^ T cells (Fig. 2A-C). Interestingly, while CD38^+^ T cells and KLRG1^+^ CD8^+^ T cells were significantly decreased, we observed a trend towards increased KLRG1^+^ CD4^+^ T cells in the spleens of aged μMT mice (Fig. S5D). These improvements within the aged T cell compartment were further observed within the blood (Fig. S5F-I), liver (Fig. S5J-M), and lung (Fig. S5N-Q), suggesting that a global absence of B cells could prevent systemic aging of the T cell compartment in multiple tissues. Ex-vivo stimulation of T cells from μMT mice confirmed decreased inflammatory potential of T cells, as CD8^+^ T cells displayed reduced IFNγ production, and CD4^+^ T cells produced limited IFNγ and IL-17, compared to WT controls (Fig. 2D, Fig. S5E). In line with these findings, aged μMT mice also possessed decreased amounts of T follicular helper (Tfh; CXCR5^+^ PD1^hi^) cells and regulatory T cells (Tregs; Foxp3^+^) (Fig. 2E-F), which represent two additional subsets of CD4^+^ T cells that are highly dysregulated within the aged spleen (*28, 29*). Surprisingly, these changes occurred in the absence of any alterations to thymic involution, thymocyte number or T cell development (Fig. 2G-H), suggesting that B cells likely regulate the T cell compartment in the periphery rather than altering thymic output or developmental pathways within the thymus.

**Fig. 2:**
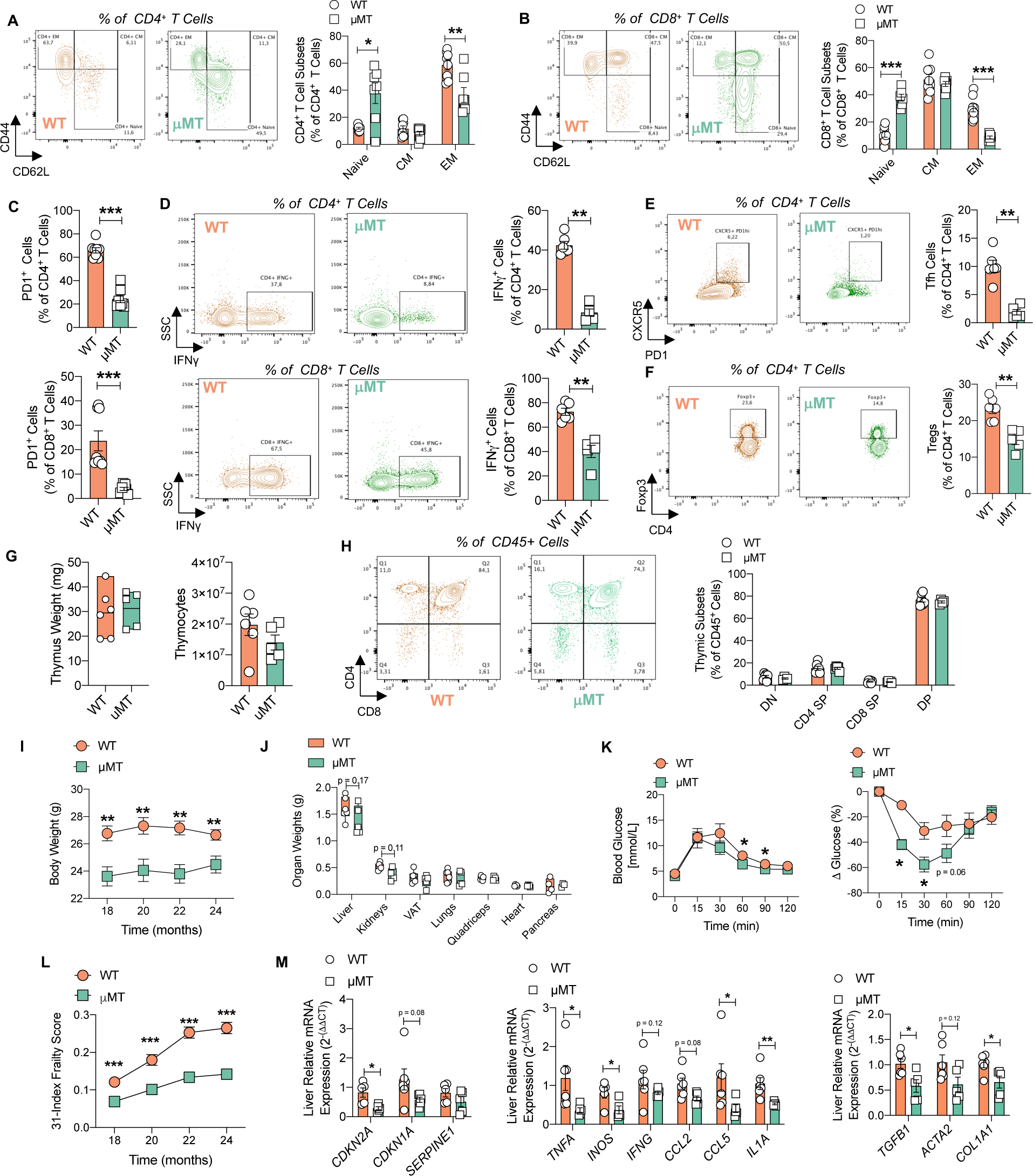
Aged µMT mice display decreased splenic T cell immunosenescence and improved age-related health parameters. (A-F) Flow cytometric assessment of T cells from spleens of aged µMT and WT mice (n=6-8 per group). (A) Representative plots (left) and relative abundance (right) of naive, effector memory and central memory in CD4^+^ T cells. (B) Representative plots (left) and relative abundance (right) of naive, effector memory and central memory cells in CD8^+^ T cells. (C) Relative abundance of PD1^+^ cells in CD4^+^ (top) and CD8^+^ (bottom) T cells. (D)Representative plots (left) and relative abundance (right) of IFNγ^+^ cells in CD4^+^ (top) and CD8^+^ (bottom) T cells post 5-hour ex vivo stimulation. (E) Representative plots (left) and relative abundance (right) of T follicular helper cells. (F) Representative plots (left) and relative abundance (right) of regulatory T cells. (G-H) Thymus and thymocyte assessment of aged µMT and WT mice (n=4-6 per group). (G) Thymus weight (left) and thymocyte yield (right). (H) Representative plots (left) and relative abundance (right) of thymic T cell subsets. (I-L) Healthspan parameters of aged µMT and WT mice (n=3-14 per group). (I) Body weight over time. (J) Organ weight. (K) Glucose tolerance test (left) and insulin tolerance test (right) results. (L) 31-index frailty scores. (M) RT-PCR gene expression of senescence (left), senescence associated secretory phenotype (middle) and fibrosis (right) markers in perfused livers. Data are means ± SEM. * denotes *p* < 0.05, ** denotes *p* < 0.01, and *** denotes *p* < 0.001.

As aged μMT mice lack B cells and harbor T cells that fail to adopt an immunosenescent phenotype, we next determined whether B cell deficiency was associated with improvements in age-related parameters of frailty and metabolic dysfunction. Compared to WT controls, aged μMT mice had a slightly lower body weight yet displayed no overt differences in organ and tissue weight at the 24-month time point (Fig. 2I-J). However, B cell deficiency was associated with improved metabolic sensitivity as determined via glucose and insulin tolerance tests (Fig. 2K), as well as improvements in a 31-index frailty score (Fig. 2L – Fig. S6A), a gold standard readout of mammalian healthspan parameters (*30*). These improvements were associated with improved organ function, as the aged μMT liver displayed decreased expression of markers linked with age-related senescence, senescence associated secretory phenotype (SASP), and fibrosis (Fig. 2M). A similar decrease in expression of SASP markers was observed in the lungs of aged μMT mice, although transcriptional markers of senescence and fibrosis remained unchanged (Fig. S6B). In line with decreased inflammatory SASP output, histological assessment with imaging scoring algorithms, confirmed significant improvements in tissue fibrosis in aged μMT mice, compared to WT controls (Fig. S6C). Cumulatively, these results suggest that an absence of B cells during the aging process is linked with a less immunosenescent T cell compartment, along with improvements in various parameters of age-associated multi-morbidity.

### B cells drive immunosenescence of the T cell transcriptional landscape and restriction of TCR diversity with age

To gain a deeper understanding of the mechanisms by which B cells direct T cell immunosenescence, we performed 5’ single cell RNA sequencing along with VDJ TCR sequencing on sorted CD3^+^ cells from the spleens of aged μMT and WT mice (Fig. 3A, Fig. S7A). A dimensional reduction of our dataset using uniform manifold approximation and projection (UMAP) revealed 16 clusters found in both conditions (Fig. 3B). Based on expression patterns of CD4 (*Cd4)*, CD8 (*Cd8a)*, CD62L *(Sell),* and CD44 (*Cd44*) we were able to annotate majority cell clusters as naive, EM and CM T cells (Fig. 3B-C). Within the aged μMT T cell pool we observed an increase in the abundance of naive CD4^+^ (cluster 3) and CD8^+^ (cluster 0) T cells (Fig. 3D). Amongst memory cells, EM CD4^+^ (cluster 1) and CD8^+^ (cluster 6) T cells were dramatically decreased in μMT mice, while the abundance of CM CD8^+^ T cells (cluster 2) was slightly higher in aged μMT mice (Fig. 3D). Naive T cells were further found to express the secondary lymphoid organ homing integrin CCR7 (*Ccr7*), while EM T cell clusters also contained cells expressing PD1 (*Pdcd1*) and Tox (*Tox)* (Fig. 3C), which have previously been thought to represent unique age-related T cell subsets (*22*). Cluster 4 corresponded to Tregs as they selectively expressed Foxp3 (Fig. S7B), and were also decreased in aged μMT mice (Fig. 3D). The remaining clusters displayed several features that prevented them from being annotated as traditional naive, EM or CM T cells. First, we identified 3 clusters of NKT cells based on their co-expression of NK1.1 (*Klrb1c*) (Fig. 3C), amongst which NKT (1) (cluster 5) and NKT (2) (cluster 11) clusters displayed an increased abundance while NKT (3) cells (cluster 14) slightly decreased in abundance (Fig. 3D); indeed, NKT cells have previously been reported to be present at a higher frequency in μMT mice and our findings are reflective of this (*31*). The remaining T cells displayed unique gene expression patterns that enabled us to annotate them based on these features: high expression of the mitochondrial enzyme leucyl-tRNA synthetase (*Lars2* – cluster 7), elevated levels of the long noncoding RNA malat1 *(Malat 1* – cluster 8), T cell receptor beta variable 3 enrichment (*Trbv3* – cluster 9), myeloid-like features (*Apoe, C1qb* – cluster 10), mixed immune cell genes (*H2-Aa, S100a9* – cluster 13), an interferon signature (*Isg15, Ifit1,Ifit3* – cluster 12), and proliferation (*mKi67, Top2a* – cluster 15) (Fig. 3B, S7B). Alterations to the abundance of these cells in aged μMT mice were minor: Trbv3^+^ naive T cells and myeloid-like CD3^+^ cells displayed slight increases, while the remaining cells displayed limited decreases (Fig. 3D). Cumulatively, these findings concurred with our previous observations made via flow cytometry analyses of T cells in aged μMT mice.

**Fig. 3:**
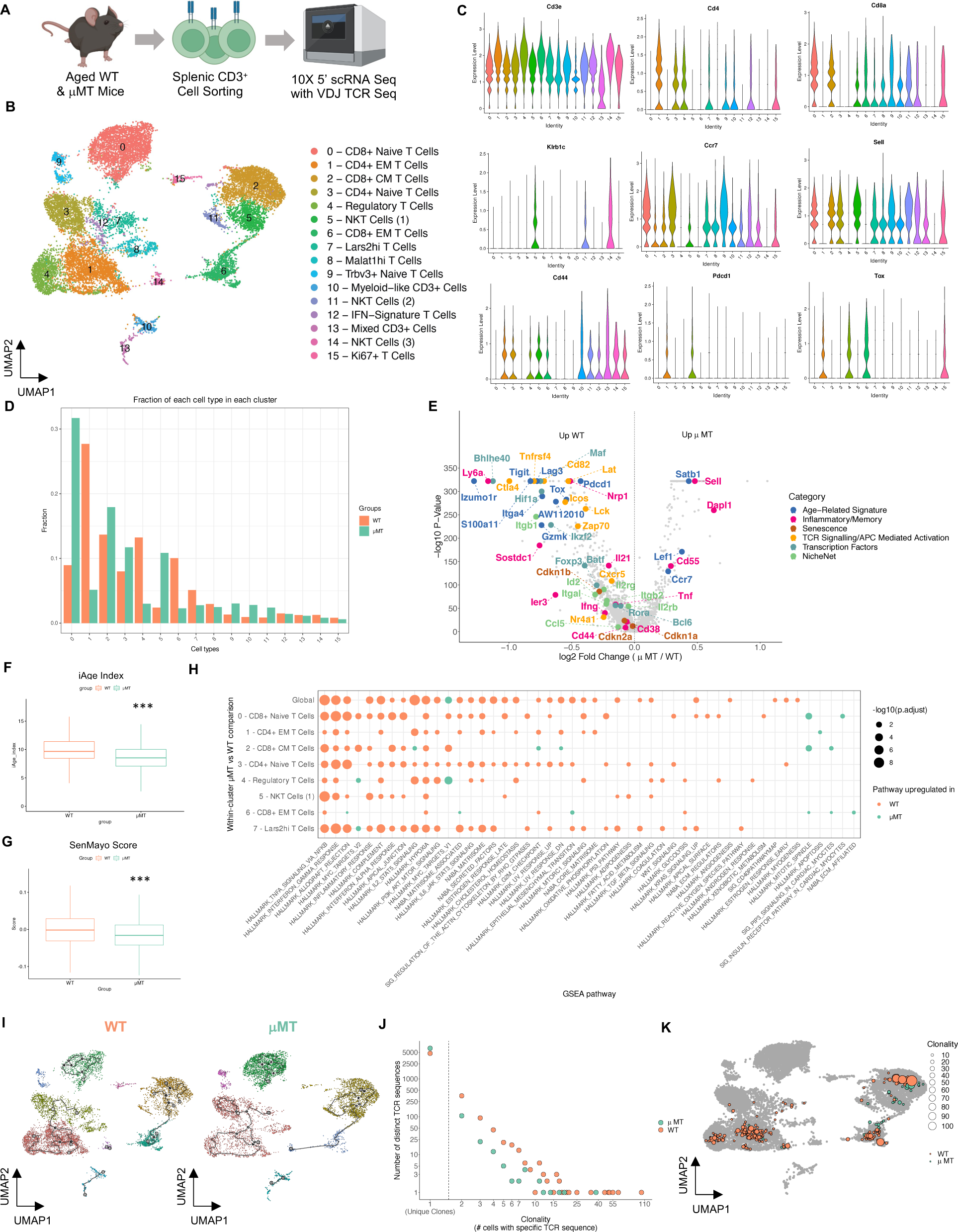
B cells shape the aging T cell transcriptional landscape and TCR repertoire. (A-K) 5’ single cell transcriptomics with VDJ T cell receptor analyses on splenic CD3^+^ cells from aged uMT and WT mice (2 pooled co-housed mice per group). (A) Schematic illustration of experiment workflow. (B) UMAP and annotation of splenic CD3^+^ clusters from aggregated aged µMT and WT mice. (C) Gene expression violin plots from aggregated aged µMT and WT mice. (D) Abundance of each cluster from µMT and WT mice. (E) Volcano plot of genes globally upregulated in CD3^+^ cells from aged µMT and WT mice. (F) iAge Index of bulk CD3^+^ cells. (G) SenMayo score of bulk CD3^+^ cells from aged µMT and WT mice. (H) GSEA pathway analysis of CD3^+^ cells from aged µMT and WT mice. (I) Trajectory analysis of CD3^+^ cells from aged µMT and WT mice. (J) TCR diversity and clonality analysis of bulk from aged µMT and WT mice. (K) TCR clonality overlaid on UMAP projection of CD3+ clusters from aged µMT and WT mice. *** denotes p < 0.001.

We next assessed differentially expressed genes (DEG) between global splenic T cells from the two conditions (Fig. 3E). WT T cells displayed upregulated expression of various identified age-related markers (*Izumo1r, Tigit, Lag3, Gzmk, S100a11, Itga4, Tox, Pdcd1, Aw112010)*, whereas μMT linked T cells displayed upregulation of transcriptional markers found commonly in young T cells (*Satb1, Lef1, Ccr7)*; these markers were identified via recent single cell transcriptomic analyses between young and aged T cells (*22, 29*). Accordingly, WT T cells displayed higher levels of traditional markers associated with inflammatory/memory T cells (*CD44, CD38, Nrp1, Ier3, Sostdc1*, *Ly6a, IL21, Tnf, Ifng)* whereas μMT T cells showed an upregulation of genes linked with naiveness and quiescence (*CD55, Sell, Dapl1)*. Importantly, in WT T cells we observed higher expression of markers linked with (1) TCR signaling and APC mediated activation (*Lck, Zap70, Icos, Cd82, Tnfrsf4, Ctla4, Cxcr5,* and *Nr4a1)*, (2) transcription factors involved with T cell differentiation (*Maf, Foxp3, Gata3, Rora, Bcl6, Batf, Ikzf2, Hif1a, Bhlhe40*), (3) T cell senescence (*Cdkn1a, Cdkn1b, Cdkn2a*), and (4) receptor/genes linked with B cell interaction, based on our previous NicheNet analyses (*Itgb1, Id2, Itgal, Itgb2, Ccl5, Il2rb, Il2rg*). At the cluster level many of these differences were observed amongst individual clusters, (Fig. S7C), suggesting that B cells can modulate the inflammatory and age-related profile of each subset and differences are not associated with shifting cell abundances.

We next applied algorithms to our datasets to enable us to explore differences between the inflammatory age (iAge index) (*25*) and senescence-associated features (SenMayo score) (*32*) of T cells from spleens of aged WT and μMT mice. Indeed, T cells from aged μMT mice had a lower iAge index (Fig. 3F, S7D) and SenMayo score (Fig. 3G, S7E) confirming their decreased immunosenescent phenotype. To gain deeper understanding of the mechanistic axes responsible for these changes we performed gene set enrichment analysis (GSEA) linked pathway analysis (Fig. 3H). Most striking was the observation that, compared to μMT T cells, WT T cells displayed higher expression of pathways linked to TNFα, IFNγ, IL-2 and TGFβ responsive signaling; these were cytokine elicited pathways that were previously identified via our NicheNet analyses to be potential axes by which B cells induce T cell dysfunction with age. Furthermore, WT T cells displayed elevations in several metabolic pathways, including oxidative phosphorylation, glycolysis, and fatty acid metabolism, that are essential for their activation and inflammatory output (*33*). Given that T cells from aged μMT mice displayed a decreased immunometabolic and immunosenescent profile, we hypothesized that in the absence of B cells, T cells would fail to undergo similar differentiation patterns that are observed in WT T cells with age. Indeed, pseudotime trajectory analyses highlighted differences in definitive fates and branching nodes in splenic T cells from aged WT and μMT mice (Fig. 3I, S7F).

Finally, given these differences to the makeup of the T cell compartment between aged WT and μMT mice, we rationalized that B cells might also be responsible for the altered T cell clonality and a loss in TCR diversity with age. Analysis of the VDJ TCR sequences unveiled that at baseline (clonality = 1) T cells from aged μMT mice displayed higher diversity and markedly less increases in clonal expansion (clonality > 1) compared to WT T cells with age (Fig. 3J). When superimposed upon our UMAP, it was clearly visible that the increase in T cell clonality with age in WT mice was linked largely with CM, EM and regulatory T cell clones, confirming that differentiation of naive cells towards these fates is responsible for a shrinking TCR repertoire (Fig. 3K). In mice lacking B cells, very few T cell clones are able to expand with age, creating a diverse TCR repertoire. As such, our findings provide direct evidence that B cells are a major cell type responsible for aging of naive T cells towards an immunosenescent fate, and for the marked TCR restriction that is seen with aging.

### CD20 monoclonal antibody therapy perturbs T cell aging

We next sought to determine whether pharmacological depletion of B cells in middle-aged mice, with a depleting CD20 monoclonal antibody (mAb), can prevent age-related T cell dysfunction. 14-15-month-old WT mice were injected with CD20 mAb or an isotype control (ISO), once every 2 weeks for 12 weeks (Fig. 4A). Upon completion of the 12-week treatment regimen, the spleen displayed a marked reduction in the relative abundance and cellularity of B cells (Fig. 4B), as expected. Accordingly, CD20 mAb treatment resulted in reductions in the B220^lo/−^ and B220^hi^ (B2) fractions, as well as B2 (B220^hi^) subsets (Fig. 4C-D). Interestingly, while we observed no differences in the relative abundance of CD11c^+^ T-bet^+^ B2 cells, their absolute cell numbers were still noticeably decreased (Fig. 4E). Our findings suggest that an acute CD20 mAb treatment regimen can successfully deplete broad splenic B cell subsets including those linked with immunosenescence.

**Fig. 4:**
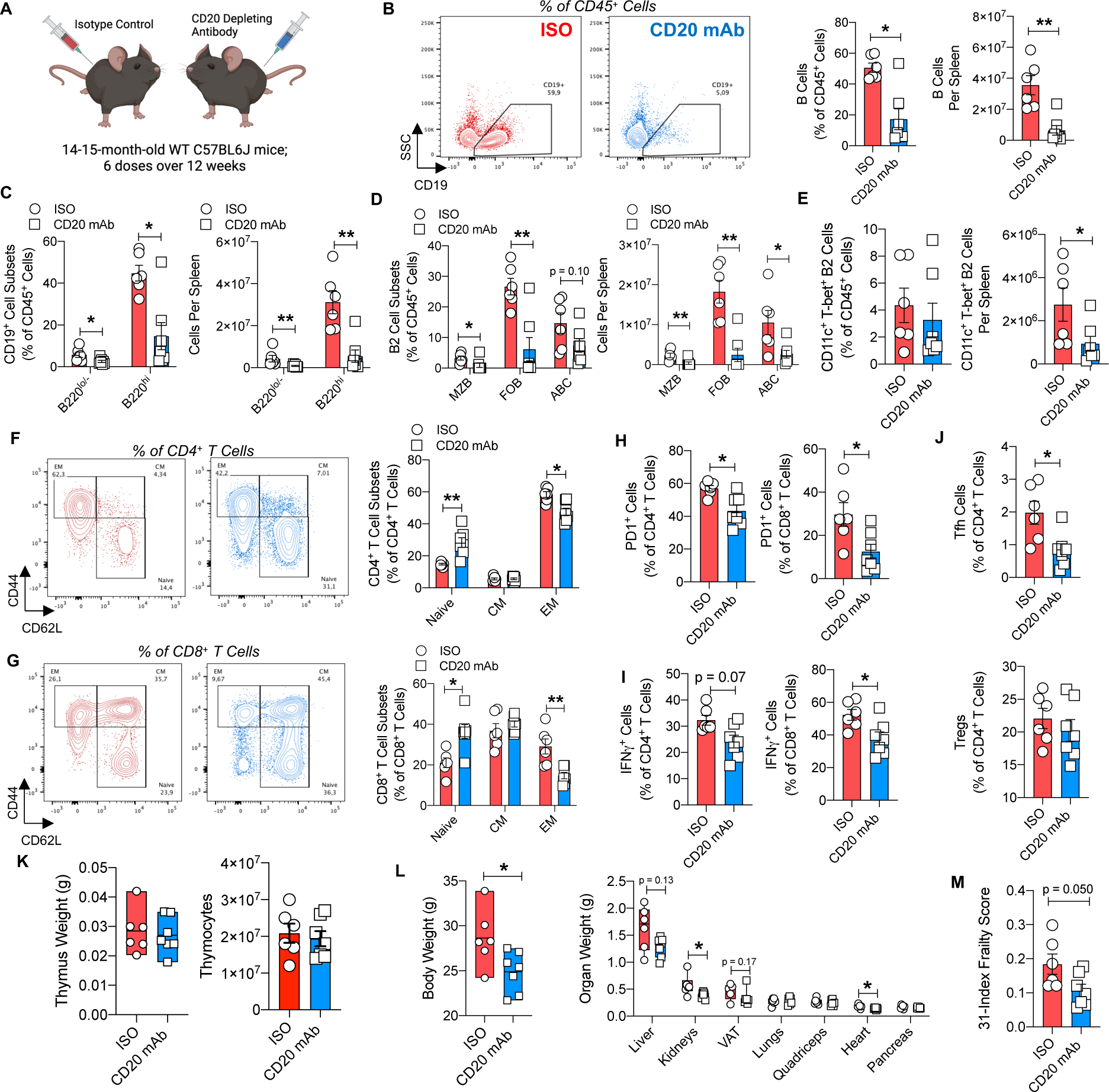
CD20 mAb therapy depletes age-related B cell subsets, halts T cell immunosenescence and improves healthspan outcomes. (A-J): Flow cytometric assessment of B and T cells from spleens of aging CD20 mAb and isotype control treated mice (n=6-7 per group). (A) Schematic illustration of experiment design. (B) Representative plot (left), relative abundance (middle) and cell number (right) of total B cells. (C) Relative abundance (left) and cell number (right) of B cell subsets. (D) Relative abundance (left) and cell number (right) of B2 cell subsets. (E) Relative abundance (left) and cell number (right) of splenic CD11c^+^ T-bet^+^ B2 cells. (F) Representative plot (left) and relative abundance (right) of naive, central memory, and effector memory subsets in CD4^+^ T cells. (G) Representative plot (left) and relative abundance (right) of naive, central memory, and effector memory subsets in CD8^+^ T cells (H) Abundance of PD1^+^ cells in CD4^+^ (left) and CD8^+^ (right) T cells. (I) Abundance of IFNγ^+^ cells in CD4^+^ (left) and CD8^+^ (right) T cells. (J) Abundance of T follicular helper cells (top) and regulatory T cells (bottom). (K) Thymus weight (left) and thymocyte yield (right) from CD20 mAb and isotype treated aged WT mice (n=6-7 per group). (L) Body weight (left) and organ weight (right) of CD20 mAb and isotype treated aged WT mice (n=6-7 per group). (M) 31-index frailty score of CD20 mAb and isotype treated aged WT mice (n=6-7 per group). Data are means ± SEM. * denotes *p* < 0.05, and ** denotes *p* < 0.01.

While CD20 mAb treatment increased splenic T cell abundance, their cellularity remained unchanged (Fig. S8A). Furthermore, treatment was associated with a mild increase in the abundance of CD4^+^ T cell fraction, while CD8^+^ T cells remained unchanged (Fig. S8B). Remarkably, WT mice treated with CD20 mAb displayed an increased abundance of naive T cells and reductions EM T cells in both CD4^+^ and CD8^+^ compartments, compared to ISO treated controls (Fig. 4F-G). Congruently, we observed decreases in the abundance of splenic PD1^+^, IFNγ^+^, and CD38^+^ T cells in CD20 mAb treated mice, while the abundance of KLRG1^+^ T cells remained unaltered (Fig. 4H-I, Fig. S8C-D). CD20 mAb treatment also decreased the expansion of Tfh cells with age, although Treg abundance remained unchanged (Fig. 4J). In line with these findings, similar observations regarding depletion of B2 cells and alterations to T cell heterogeneity were made in the blood (Fig. S8E-H), liver (Fig. S8I-L), and lung (Fig. S8M-P), positing potential systemic therapeutic efficacy of CD20 mAb treatment. Importantly these changes to the T cell compartment were once again observed in the absence of any changes to thymic size or thymocyte number, confirming that B cells utilize thymic independent mechanisms to modulate the T cell compartment with age (Fig. 4K).

CD20 mAb treatment in aged mice has previously demonstrated therapeutic efficacy in preventing age-associated metabolic dysfunction, which is directly linked with the activity of adaptive immune cells (*34*). Consequently, we speculated that CD20 mAb treated mice might also display improvements in the 31-index frailty score and health condition. Indeed, CD20 mAb treated mice possess a lower body weight, which was associated with significant reductions in the weight of the kidneys and heart (Fig. 4L). Furthermore, pre-sacrifice assessment revealed a strong trend towards improved frailty outcomes in CD20 mAb treated mice, compared to ISO controls (Fig. 4M). These changes were likely linked to an improvement in overall health with trending improvements seen across multiple measures, as the only statistically significant individual improved measure was a decrease in piloerection (Fig. S9A). Cumulatively, these findings suggest to us that B cells can be actively targeted in late adulthood to prevent age-related decline in the naive T cell compartment and mammalian healthspan.

### Insulin facilitates aged B cell responses

Given that our findings and previous reports suggest a shift towards a pro-inflammatory tone in B cells during the aging process, we next sought to determine mechanisms fueling their pathogenic phenotype with age. To meet the higher energy demands associated with a pro-inflammatory output, immune cells undergo shifts in cellular metabolism (*35*). Recent evidence suggests that aged B cells display a hypermetabolic profile linked with their pro-inflammatory capacity, a phenotype that is more pronounced in ABCs compared to FOBs (*11*). Despite this understanding, internal triggers that contribute to the cellular metabolism of B cells with age remain relatively unknown.

The insulin signaling pathway downstream of the insulin receptor (InsR) is an evolutionary conserved signaling axis that boosts cellular metabolism and is negatively correlated with longevity across species (*36*). Cells of the adaptive immune compartment possess the InsR and upregulate its expression upon activation (*37*). Particularly for T cells, InsR signaling facilitates changes in metabolism required for their inflammatory output necessary for viral clearance and autoimmune responses (*38*). Yet whether the same metabolic axes in B cells are evoked is unknown. Given that aging is associated with hyperinsulinemia, elevated B cell metabolism, and increased B cell effector function, we investigated whether insulin may serve as an environmental cue that harmonizes these observations and whether it is responsible for driving age-related B cell dysfunction.

To test this hypothesis, we further mined our previously utilized public data set comparing systemic young and aged B cells (*22*), and investigated if insulin-related pathways enriched among genes with higher expression in aged B cells (Fig. 5A). Indeed, aged B cells displayed an increased GeneRatio score in pathways related to the phosphatidyl inositol-kinase (PI3K)/protein kinase B (AKT) signaling axis driving the activation of the mammalian target of rapamycin (mTOR) (Fig. 5A). Phosphorylation of Akt at sites Thr308 and Ser473 are key events required for the activation of mTOR resulting in a cascade of events necessary for metabolic reprogramming of B cells (*39*). Indeed, at the molecular level, B2 cells from aged spleens displayed greater basal phosphorylation at the Thr308 site and a trend towards increase at the Ser473 site, compared to young B cells (Fig. 5B). Evaluation of aged B2 cell subsets unveiled that in contrast to the majority FOB population, CD11c^+^ T-bet^+^ ABCs displayed elevated basal phosphorylation of Akt at both sites (Fig. 5C) positing that they do possess greater metabolic demands, in line with previous findings (*11*).

**Fig. 5.**
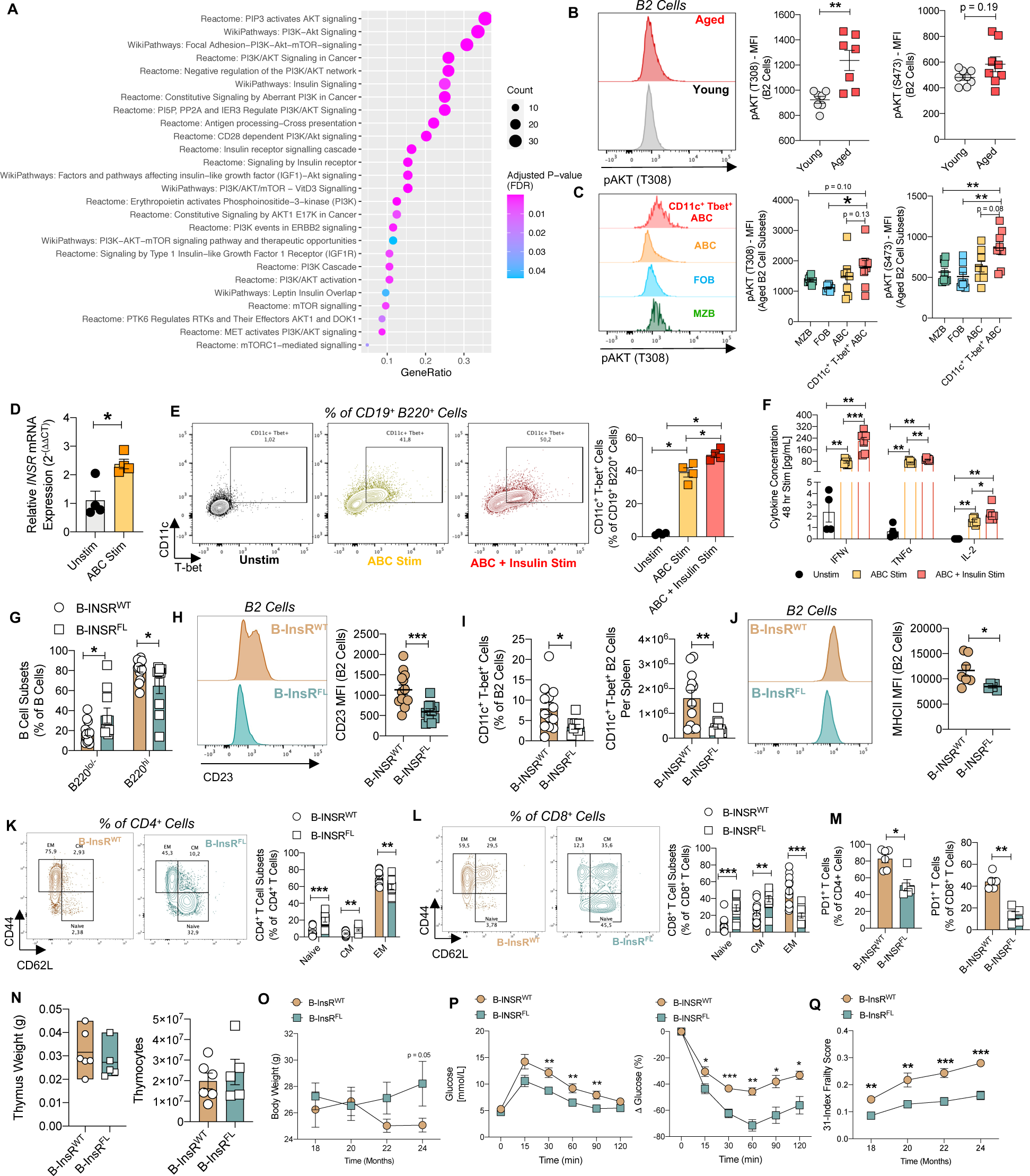
Insulin boosts age-related B cell responses linked with an immunosenescent T cell compartment. (A) GeneRatio analysis of bulk splenic, hepatic, pulmonary, and peritoneal B cells highlighting insulin signaling related pathways upregulated in aged WT mice compared to young WT mice (Mogilenko *et al.* GSE155006). (B) Representative plot of pAKT-T308 (left), MFI of pAKT-T308 (middle) and MFI of pAKT-S473 (right) in young and aged splenic B2 cells (n=7-8 per group). (C) Representative plot of pAKT-T308 (left), MFI of pAKT-T308 (middle) and MFI of pAKT-S473 (right) aged splenic B2 cell subsets (n=8). (D) Expression of *INSR* mRNA in unstimulated and ABC-stimulated B cells (n=4 per group). (E)Representative plots (left) and abundance (right) of CD11c^+^ T-bet^+^ cells from unstimulated, ABC stimulated, and ABC + Insulin stimulated B cells (n=4 per group). (F) Culture supernatant cytokine concentration from unstimulated, ABC stimulated, and ABC + Insulin stimulated B cells (n = 4-7 per group). (G-M) Flow cytometric assessment of B and T cells from spleens of aged B-InsR^WT^ and B-InsR^FL^ mice (n=5-13 per group). (G) Relative abundance of B cell subsets. (H) Representative plot (left) and MFI (right) of CD23 expression in B2 cells. (I) Abundance (left) and cell numbers (right) of CD11c^+^ T-bet^+^ B2 cells. (J) Representative plot (left) and MFI (right) of MHCII expression in B2 cells. (K) Representative plot (left) and relative abundance (right) of naive, central memory, and effector memory subsets in CD4^+^ T cells. (L) Representative plot (left) and relative abundance (right) of naive, central memory, and effector memory subsets in CD8^+^ T cells. (M) Abundance of PD1^+^ cells in CD4^+^ (left) and CD8^+^ (right) T cells. (N) Thymus weight (left) and thymocyte yield (right) from aged B-InsR^WT^ and B-InsR^FL^ mice (n=5-6 per group). (O-Q) Healthspan parameters of aged B-InsR^WT^ and B-InsR^FL^ (n=5-14, per group). (O) Body weight over time. (P) Glucose tolerance test (left) and insulin tolerance test (right) results. (Q) 31-index frailty scores over time. Data are means ± SEM. * denotes *p* < 0.05, ** denotes *p* < 0.01, and *** denotes *p* < 0.001.

The formation of CD11c^+^ T-bet^+^ B cells has been identified to be dependent upon a combination of stimuli, including toll-like receptor (TLR), B cell receptor (BCR) and IL-21/IFNγ cytokine signals (*14*). Remarkably, *in vitro* stimulation of B cells with these triggers (ABC stim) resulted in increased transcriptional expression of the InsR (Fig. 5D). Furthermore, supplementing ABC stimulated B cells with insulin expanded the *in vitro* generation of CD11c^+^ T-bet^+^ B cells (Fig. 5E) and boosted their production of T cell modulating pro-inflammatory cytokines (IFNγ, TNFα, IL-2) (Fig. 5F). Thus, B cell activation and subsequent upregulation of the InsR are perhaps reflective of a greater need for insulin consumption, which acts as a metabolic adjuvant via the Akt signaling pathway to support the generation of CD11c^+^ T-bet^+^ ABCs, and their downstream cytokine production abilities necessary for T cell modulation.

### Lack of B cell specific InsR attenuates age-related adaptive immune dysfunction and disease outcomes

To formally test the role of InsR signaling in age-related pathogenicity of B cells, we generated and aged mutant mice within which B cells specifically lacked the InsR (CD19 Cre^+/−^ InsR^FL/FL^; B-InsR^FL^ mice) and compared them against age-matched wild type Cre controls (CD19 Cre^+/−^ InsR^WT/WT^; B-InsR^WT^ mice). We first surveyed the effects of InsR ablation on aged splenic B cells and found no alterations to bulk B cell abundance and cellularity (Fig. S10A). However, within the B cell compartment, we observed a marginal decrease in the abundance of B2 cells and a concurrent increase in the abundance of B220^lo/−^ cells (Fig. 5G). Of note, InsR deficient B2 cells failed to express typical levels of CD23 by flow cytometry (Fig. 5H), which prevented us from defining aged B2 cell subsets based on expression of CD21 and CD23 (Fig. S10B). However, in line with our *in vitro* findings, B-InsR^FL^ mice displayed a decrease in abundance and cellularity of CD11c^+^ T-bet^+^ B2 cells (Fig. 5I). Furthermore, while CSR remained unchanged in InsR deficient B2 cells (Fig. S10C), we noted a decrease in their expression of MHCII (Fig. 5J), positing a potential decline in their ability to stimulate T cells. As such, we further speculated that aged B-InsR^FL^ mice would possess an altered splenic T cell compartment, compared to age-matched B-InsR^WT^ control mice.

While the abundance and cellularity of bulk T cells remained unchanged, we observed a decrease in the abundance of CD4^+^ T cells and an increased abundance of CD8^+^ T cells in aged B-InsR^FL^ mice (Fig. S10D-E). Remarkably, within both CD4^+^ and CD8^+^ T cell compartments we observed increased abundances of naive T cells and reductions in the abundance of EM T cells in the spleens of aged B-InsR^FL^ mice (Fig. 5K-L). These findings were further accompanied with a significant decrease in the abundance of PD1^+^ T cells (Fig. 5M), and a trend towards decreased IFNγ secreting T cells and Tfh cells, in the spleen, while Tregs remained unchanged (Fig. S10F-G). Similar changes in memory and naive T cells were observed amongst hepatic T cell subsets and pulmonary CD8^+^ T cells in aged B-InsR^FL^ mice (Fig. S10H-M), and cumulatively, these findings were once more observed to be independent of any changes to thymic involution or thymocyte number (Fig. 5N).

We next assessed parameters of age-related decline in aged B-InsR^FL^ mice compared to B-InsR^WT^ controls. Despite no changes to body weight or organ weight (Fig. 5O, Fig. S10N), aged B-InsR^FL^ mice displayed significant improvements in glucose and insulin tolerance (Fig. 5P), findings that are in line with a restored adaptive immune compartment. Improvements in metabolic homeostasis of aged B-InsR^FL^ were further associated with decreased scores on the 31-index frailty index (Fig. 5Q, Fig. S10O), thereby suggesting improvements in overall healthspan, compared to age-matched B-InsR^WT^ controls. Our findings indicate that InsR signaling in B cells is a driving pathway for B cell induced adaptive immunosenescence and age-related decline of various health parameters.

## Discussion

During aging, one of the most well documented changes to the immune system is the pronounced loss in naive T cells and associated loss in TCR diversity. Age-related T cell subsets assume non-redundant roles in their control over inflammaging, yet collectively impact the T cell repertoire due to clonal expansion of activated T cell subsets. The loss in TCR diversity with age is a major mechanism of immune system vulnerability to new infections, such as COVID-19 (*7*), and to cancer in the elderly. While the dominant mechanisms influencing the loss of naive T cells with age are not completely understood, it is thought to occur through a combination of expansion of memory, senescent, and exhausted T cell subsets, as well as potential clonal transformation of T cells (*9*). Other hypotheses postulating the mechanisms driving this phenomenon highlight the role of thymic involution, metabolic defects, and epigenetic alterations (*8*), yet whether there are dominant players responsible for the compositional changes to the aged T cell repertoire has been understudied. Aging is also associated with a concurrent dysregulation of the B cell compartment, which is most noticeably demarcated by the expansion of an antigen-experienced ABC subset that also harbors CD11c^+^ T-bet^+^ B cells, yet little is known about the consequences to these alterations.

Here, we find that B cells engage in age-related crosstalk with T cells and support the phenotypic aging of the T cell compartment across the lifespan of mice. We demonstrate that B cell mediated activation of T cells results in transcriptional changes that evokes their differentiation towards pathogenic states consisting primarily of memory T cell subsets. Cumulatively, B cell differentiated T cells display a higher propensity towards an inflammatory and senescent phenotype, which directly impacts TCR diversity and drives the clonal expansion of T cells with age. Our findings complement previous reports suggesting that an aged B cell compartment displays superior T cell antigen presenting capabilities (*40*). Indeed, compared to FOBs, ABC-associated CD11c^+^ T-bet^+^ B cells express higher levels of T cell stimulatory machinery, and form longer and more stable interactions with T cells, resulting in greater stimulation and proliferation of T cells (*40*). Similarly, superior antigen presentation abilities of CD11c^+^ T-bet^+^ B cells has also been linked to the activation and differentiation of T cells in mouse models of autoimmunity (*40, 41*).

Intercellular communication and pathway analysis algorithms cumulatively highlighted mediators utilized by B cells to exert their pathogenicity, consisting of a combination of secreted and surface factors. Among these mechanistic axes, instigation of T cell inflammatory cytokine receptor signaling nodes were most prominent. Indeed, ABCs have been noted to display elevated inflammatory cytokine output (*11, 42, 43*), but the consequences of these changes across the lifespan of mice was largely unknown. Our findings suggest that the adoption of a pro-inflammatory phenotype by B cells likely supports the generation and maintenance of age-related T cell populations, thereby driving T cell immunosenescence and features of aging. Given this role for B cell crosstalk onto T cells in driving adaptive immunosenescence, it will also be important to determine whether MHC complexes of B cells harbor unique peptide sequences that are linked with an aged environment. As such, an assessment of the B cell MHC peptide pool might provide valuable insights into specific antigens driving T cell dysfunction with age, thereby suggesting that aging is associated with antigen specific, including potentially infectious or auto-immune like processes (*44*). Furthermore, we observed dysregulation of splenic microanatomical structure as well as the formation of tertiary lymphoid structures with age, likely prompting increased physical interaction between B cells and T cells. Indeed, ABC-associated CD11c^+^ T-bet^+^ B cells have been found to naturally localize at the splenic T cell/B cell border (*40*), and ectopic lymphoid cluster formation has been documented previously within the aged liver (*45*) and visceral adipose tissue (*34*). However, the importance of spatial proximity between aged B cells and T cells, and whether it is essential in facilitating crosstalk, remains to be determined. Finally, while we noted concurrent alterations to B cells and T cells within the spleen, liver, lung and blood, there are many additional non-lymphoid organs that warrant further examination. For instance, the adipose tissue (*34*) and meninges (*46*) are two additional sites where B cells display age-related perturbations to their phenotype and possess features that distinguish them from their splenic counterparts. As such, a more thorough investigation of B cells in tissue specific niches is warranted, and it remains to be determined whether they evoke additional mechanisms of T cell immunosenescence.

Accordingly, our models of B cell deficiency highlight the role of B cells in influencing overall decline in mammalian health with age. Indeed, aged B cell deficient mice displayed improvements in metabolic parameters, senescence, fibrosis, and frailty outcomes, implying that B cells negatively impact mammalian healthspan. CD20 mAb therapy has previously been demonstrated to bestow metabolic improvements in aged mice, notably by restoring glucose homeostasis, insulin sensitivity and adipose tissue lipolysis (*34, 47*). Here we expand upon the mode of action of CD20 mAb therapy and demonstrate that targeted depletion of B cells, given in late adulthood in mice, can halt the aging of the systemic T cell compartment, most notably preventing the formation of EM T cells and conserving the naive T cell pool, and is associated with improved age-related frailty outcomes with age. Although, it is important to note that CD20 mAb treatment does not deplete terminally differentiated antibody secreting cells, which accumulate with age (*13*) and also produce pro-inflammatory cytokines, particularly in the bone-marrow (*48*). Importantly the bone marrow is a niche for memory T cell survival and quiescence (*49, 50*), and it remains to be determined whether B cell depletion therapies can modulate this pool of T cells. Furthermore, while a dysfunctional T cell compartment is likely sufficient to drive these changes, we speculate that the age-related B cell pro-inflammatory cytokine secretion, antibody production, and T cell modulation act in concert to drive age-associated multi-morbidity.

Due to their aberrant formation and potential pathogenic function, mechanisms that result in the formation of ABCs and CD11c^+^ T-bet^+^ B cells are of broad interest. In particular, CD11c^+^ T-bet^+^ B cells have previously been demonstrated to form as a result of nucleic acid sensing via TLR7/9, ligation of the BCR, and IL-21 and IFNγ cytokine signaling (*14*). Here, we identify insulin as an additional environmental trigger that supports the formation of CD11c^+^ T-bet^+^ B cells. As ABCs have greater metabolic needs it is likely that insulin receptor signaling is required for them to undergo metabolic processes, such as glycolysis, to support their expansion and pro-inflammatory cytokine output. Indeed, a similar function of the InsR has been observed on T cells, whereby InsR signaling promotes T cell inflammatory output by boosting their cellular metabolism (*38*). Accordingly, ablation of InsR from B cells diminishes the formation of CD11c^+^ T-bet^+^ B cells and their expression of MHCII, thereby preventing age-related T cell immunosenescence. Interestingly, we find that an absence of B cells or knockout of their InsR also results in a decline in IFNγ^+.^T cells, as well as IL-21 secreting PD1^+^ CD4^+^ T cells, suggesting that the presence of aged B cells likely forms a positive feedback loop whereby T cell derived cytokines continue to support CD11c^+^ T-bet^+^ B cell expansion in congruence with TLR7/9, BCR and InsR signals. Interestingly, improvements in age-associated metabolic disease parameters were observed in mice within which Tregs specifically lacked the InsR (*51*), positing that insulin may exert its anti-longevity effects via multiple immune cells and not just B cells. Recently, the InsR was noted to associate with nuclear RNA polymerase II at promoter sites to modulate gene regulation of pathways linked with inflammation and antigen specific responses (*52*). Whether this function of the InsR is conserved in aged B cells, or other immune cells, remains to be determined but could be a potential promising avenue of future research.

Collectively, our results support a model where intrinsic age-associated environmental triggers, including InsR signaling, drive the aging of the B cell compartment. Aged B cells thereby contribute to inflammaging, by supporting T cell immunosenescence, which results in a loss of the naive T cell pool and TCR diversity, and facilitates the formation of a clonally restricted inflammatory T cell compartment. These results place B cells as key orchestrators of adaptive immune system aging, with the capacity to control mammalian healthspan, and suggest new immunotherapeutic approaches to curb the aging process.

## Acknowledgements

We thank Princess Margaret Genomics Centre (PMGC) for their help in single cell sequencing and the Princess Margaret Hospital (PMH) flow core for their assistance with flow cytometry. We would like to thank Helena Lei for help early on in mouse colony maintenance. Graphical illustrations were made via BioRender.com. This work was supported in part through funds derived from the National Institutes of Health (NIH) grant R01DK128435 (D.A.W.), and the Canadian Institutes of Health Research (CIHR) grants FDN-148385, PJT169175 and PJT186165 (D.A.W.). I.J. was supported in part by funding from Natural Sciences Research Council (NSERC #203475), Canada Foundation for Innovation (CFI #225404, #30865), Ontario Research Fund (RDI #34876), IBM and Ian Lawson van Toch Fund. This project was also supported by funds derived from an Impetus Longevity Grant from Norn Group, awarded to S.K. S.K. is also a recipient of the Queen Elizabeth II Graduate Scholarship in Science and Technology (QEII-GSST)/Aventis Pasteur, and the Dick and Peggy Sharpe Student Fellowship in Immunology.

## Author Contributions

S.K., M.C., S.W. and D.A.W. contributed to the experimental design, execution and analysis, and writing of the manuscript of this study. N.C., T.W., Y.T.C., F.J.A., assisted with *in vitro* and *in vivo* experiment execution. F.W., A.S., J.D.S., and M.K. performed computational analyses of single cell datasets. K.N., and Y.H. performed analyses of the 1000 immunomes dataset. M.W., R.K.L., M.H., I.J., A.J.G., P.S.O., D.F., S.T., and S.W. supervised aspects of the study. D.A.W. supervised the overall implementation of the study. All authors had an opportunity to view and edit the manuscript.

