## Supplementary Materials for "B Cells Promote T Cell Immunosenescence and Mammalian Aging Parameters"

This PDF file includes:

- Materials and Methods
- Table S1
- Figs S1-S10

### Materials and Methods

#### Mice

C57BL/6J (000664), CD19-cre (006785), $Insr^{fl/fl}$ (006955), $\mu$MT mice (002288) were purchased from Jackson Laboratory. CD19 $cre^{+/-}$ $Insr^{fl/fl}$ mice were generated in-house by inter crossing CD19-cre mice with $Insr^{fl/fl}$ mice. Mice were maintained in a pathogen-free, temperature-controlled, and 12 h light and dark cycle environment at the Toronto Medical Discovery Tower animal research facility. Female mice were used for experiments, unless otherwise specified. All the experimental procedures were performed under the approval of Animal User Protocol by the Animal Care Committee at the University Health Network.

#### Immune Cell Isolation

Mice were euthanized using $CO_2$ fixation prior to collecting organs, blood, and lymphoid organs. Spleens were processed into single cell suspensions and filtered through 40 $\mu$m cell strainer followed by hemolysis. Blood was collected in heparin (1000u/ml, BioShop) by cardiac puncture after euthanization or saphenous vein from live mouse and subjected to hemolysis for single cell suspension. Thymus were processed into single cell suspensions and filtered through 40 $\mu$m cell strainer without hemolysis. Livers were perfused and dissociated using gentleMACS Dissociator (Miltenyi Biotech), as previously described (*53*). Lungs were perfused and digested with collagenase IV (1 mg/ml, Sigma) for 1 h at 37°c. Single cell suspensions were isolated from lung and liver using percoll density gradient centrifugation followed by hemolysis.

#### Flow Cytometry

Single cell suspensions from different mouse organs and blood were stained with Zombie ultraviolet dye (Biolegend) for viability at room temperature for 20 min followed by Fc-blocking with CD16/32 (93, Biolegend) at 4°c for 20 min. For staining of intracellular cytokines, single cell suspensions were stimulated with phorbol myristate acetate in the presence of Golgi Stop (eBioscience) for 5 hours prior to surface staining. The cells were further stained with fluorophore-conjugated antibodies for 30 min at 4°C in dark using CD45 (30-F11), CD3 (17A2), NK1.1 (PK136), CD4 (RM4-5), CD8 (53-6.7), CD44 (IM7), CD62L (MEL-14), PD-1 (29F.1A12), KLRG1 (2F1), CD38 (90), CD19 (6D5), B220 (RA3-6B2), CD21 (7E9), CD23 (B3B4), CD11c (N418), IgM (RMM-1), IgD (11-26C.2A), CD80 (16-10A1), PD-L2 (TY25), I-Ab MHC Class II (AF6-120.1), CXCR5 (L138D7), and CD5 (53-7.3) (Biolegend). Intracellular staining was done for Tbet (4B10), IL-17 (TC11-18H10.1), IFNg (XMG1.2), and Foxp3 (150D) for 45 min at RT in dark following fixation and permeabilization using Foxp3 staining buffer set (eBioscience). Phospho-flow for pAKT staining was performed as previously described (*54*). Briefly, cells were stained with Zombie ultraviolet dye (Biolegend) for viability at room temperature for 2 min followed by Fc-blocking with CD16/32 (93, Biolegend) at 4°c for 2 min. Surface markers were stained for 7 min at 4°C. Intracellular staining was done for pAKT-Ser473 (SDRNR, Invitrogen), pAKT-pT308 (J1-223.371, BD Biosciences) for 45 min at RT in dark following fixation and permeabilization overnight at 4°C using Foxp3 staining buffer set (eBioscience). Cells were analyzed on LSRFortessa X-20 at the Princess Margaret Hospital Flow Cytometry Facility, University Health Network (Toronto, Canada). All the raw data for flow cytometry was analysed using FlowJo (v.10.7.1).

**Gene Expression Assays**
Total RNA was extracted from flash frozen cell pellets and organs using RNeasy Mini Kit (QIAGEN). Reverse–transcription was performed using SensiFAST cDNA Synthesis Kit (Bioline). qPCR was performed on QuantStudio 6 Flex Real-Time PCR system (Thermo Fischer) using SYBR Green Master Mix reagent (Applied Biosystems). Samples were normalized to housekeeping gene HPRT, GAPDH or β-actin. Relative changes in gene expression were calculated based on the ΔΔCT method using the equation $2^{-\Delta\Delta CT}$. Fold changes were presented in comparison to the control groups. See Table S1 for primer sequences.

**ELISA**
Cytokine concentration for IL2, IFNγ, and TNFα was measured in supernatant using kits (Biolegend) following manufacturers protocol.

***In Vitro* ABC Generation**
Spleens were processed into single cell suspensions and filtered through 40 $\mu$m cell strainer followed by hemolysis. B cells were purified from single cell suspensions using Pan B cell Isolation kit (Miltenyi Biotech) and 200,000 cells/well were plated in 96-well, U-bottom plate (Falcon) cultured in 200 $\mu$l RPMI-1640 (Wisent) supplemented with Penicillin (100 U/ml, Sigma), Streptomycin (100mg/ml, Sigma), L –glutamine (2 mM, Sigma), 2-mercaptoethanol (50 mM, Sigma), and 10% heat-inactivated FBS (Wisent) with or without: R848 (500 ng/mL, Miltenyi Biotech); anti-mouse IgM F(ab')$_2$ fragment (1 μg/mL, Biolegend); IL-21 (50 ng/mL, Biolegend); Insulin (100 ng/ml). B cells were cultured for 48 hours before collection of supernatant and cell pellet.

**Histology**
Tissue pieces were fixed in 10% formalin and were assessed for collagen content after sectioning using Masson's trichrome staining done by Pathology Research Program at University Health Network (Toronto, Canada). The percent collagen area was assessed on images taken at 10x resolution on 10 different LPF/section for liver, kidney, and heart and at 5x resolution on 2 different LPF/section for lung in each mouse. Blue staining of collagen was segmented from different colors using Weka Trainable Segmentation plugin classifier in FIJI software (ImageJ, v.2.3.0/1.53q). The segmented color of collagen was isolated using thresholding, keeping the thresholding limits constant across all images. The segmented image of isolated color of collagen was converted into a binary file to analyse the percent area positive for collagen (*53*).

**Anti CD20 mAb Treatment**
14-15 months old C57BL/6J mice were injected intraperitoneally with anti-CD20 monoclonal antibody (300$\mu$g, SA271G2, Biolegend) or Rat IgG2b, isotype control antibody (300$\mu$g, RTK4530, Biolegend), once every 2 weeks for 12 weeks.

**Metabolic Tolerance Tests**
Metabolic tests were conducted as previously described (*55*). Mice were fasted overnight for GTTs and for 6-7 hours in the morning for ITTs. GTTs and ITTs were performed using 1.5 g glucose per kg body mass and 0.5 U human insulin per kg body mass via intraperitoneal injection. Blood glucose measurements were made at 0-, 15-, 30-, 60-, 90-, and 120-minute time-points with a glucometer (Counter next *EZ*).

### Frailty Scoring

Health span of mice was assessed using a 31-item frailty index to identify humane interventions and endpoints. The scoring was done every 2 months, starting at 18 months. The 31-item frailty index is a list of non-invasive clinical assessment of 31 potential deficits in aging mice that are scored 0, 0.5 or 1 based on the severity which generates an average score for each mouse (*30*).

### Immunofluorescence

Tissues were fixed with 4% PFA (pH 7.4) for 4h, cryo-protected with 30% sucrose solution overnight, and embedded in OCT medium. 5-10μm-thick sections were obtained, washed with PBS, and blocked for 15min with 5% Goat Serum and 0.1% Triton-X in PBS. Sections were incubated with fluorophore-conjugated primary antibody overnight at 4°C, washed with PBS, and counter-stained with DAPI. Sections were mounted with Prolong Gold anti-fade mounting medium (ThermoFisher Scientific) and imaged on an Axio Imager fluorescent microscope (ZEISS).

### T Cell Single Cell Analyses

#### Cell Sorting and Preparation

Cells from spleens of aged $\mu$MT and WT mice were prepared as described above. Cells were blocked with Fc-block CD16/32 (93) and stained with viability dye zombie aqua, CD45 (30-F11), CD3 (17A2), CD19 (6D5) (BioLegend), using methods described above for surface staining. Live $CD45^{+}$ $CD3^{+}$ $CD19^{-}$ were sorted using a MoFlo Astrios Sorting Machine, at the Princess Margaret Hospital Flow Cytometry Facility, University Health Network (Toronto, Canada), and collected into Eppendorf tubes containing a 500 uL cushion of RPMI-1640 (Wisent) with 10% heat-inactivated FBS (Wisent). Samples were processed according to protocols for 5' v2 chemistry with VDJ (T) library enrichment from 10X Genomics at the Princess Margaret Genomics Centre and sequenced on a NovaSeq 6000 instrument. Post sequencing, to align read and generate feature barcode matrices raw data was subjected to the Cell Ranger pipeline -6.1.2. Cell Ranger outputs indicated the following yields: WT CD3+ cells 8,999 cells with 7,405 cells with productive VJ spanning pairs and $\mu$MT CD3+ cells 8,312 cells with 6,268 cells with productive VJ spanning pairs.

#### Processing:

All analyses were performed in R (v.4.2.0), primarily using the Seurat package (v.4.1.1) and custom analysis scripts (*56, 57*). Single-cell RNA and T-Cell Receptor (TCR) sequencing data matrices resulting from the 10x CellRanger (10x Genomics) software pipeline were loaded, formatted, and merged. Exploratory analyses examining standard quality control metrics were used to verify data quality, resulting in a removal of cells containing >10% mitochondrial RNA and <250 genes/features. Doublet cells were identified and removed from the downstream analysis by using the DoubletFinder (v.2.0.3) R package (*58*) with parameters PCs=1:20, pN=0.25, and nExp=7.5%. Raw RNA counts were first normalized and stabilized with the SCTransform v2 function (SCT), then followed by the reciprocal PCA (RPCA) integration workflow for joint analysis of two single-cell datasets. In doing so, the top 3,000 highly variable genes/features among the datasets were used to run SCT; and then 2,000 highly variable genes/features and the 20 top principal components (PCs) with k.anchor=4 were used to find "anchors" for integration.

To identify putative cell subsets (clusters), the 20 top principal components (PCs) summarizing the RNA expression of each cell were used to perform Seurat clustering with a resolution parameter of 0.5, resulting in 16 RNA clusters each containing a mix of cells from the WT and $\mu$MT samples. RNA differential expression ("Marker") analyses were used to find up- and down-regulated RNAs for each Seurat cluster (vs. all other clusters), between all $\mu$MT and WT cells regardless of the cluster, and RNA differences between $\mu$MT and WT cells within each of the 8 largest Seurat clusters (clusters 0 through 7). Smaller clusters were omitted due to few cells per cluster. In all cases, Seurat FindAllMarkers/FindMarkers analyses were run with no log-fold-change pre-filtering (logfc.threshold=0) using the MAST algorithm (*59*) .

<u>*Pathway Analysis:*</u>
Following differential expression, Gene Set Expression Analysis was used to discover higher-level cellular pathways dysregulated in each comparison (*60*). Genes with p-values <=0.05 were pre-ranked by their Log2FC and incorporated into the Gene Set Enrichment Analysis (GSEA) below. Geneset collections from the Molecular Signatures Database (MSigDB v.7.5.1), specifically the Hallmark and Canonical Pathways collections, were used to perform GSEA.

<u>*Trajectory Analysis:*</u>
To study the inferred trajectory of T cell differentiation, cell trajectory analysis was performed on WT and $\mu$MT samples, respectively, by using the R package Monocle 3 (v.1.2.9) (*61, 62*). We first subset Seurat data to WT and $\mu$MT groups then run the functions as.cell_data_set(), cluster_cells(), and learn_graph(). Finally, run order_cells() with the selection of primary naive T cells as the root of the trajectory.

<u>*TCR Clonality Analysis*</u>:
The Wilcoxon Rank-Sum test and Kolmogorov-Smirnov test were used to assess whether the distributions of clonally-expanded cell populations between the WT and $\mu$MT cell populations were significantly different. In both cases, the distributions were deemed significantly different with a very low p-value ($p < 2.2 \times 10^{-16}$), with the WT cells exhibiting greater clonal expansion across larger numbers of unique TCR clones.

<u>*iAge Index and SenMayo Score Calculation:*</u>
The inflammatory aging (iAge) index was calculated by multiplying normalized and scaled gene expression with the corresponding coefficient of the gene in the iAge gene set (*25*). Cellular senescence was scored using AddModuleScore function in Seurat with the SenMayo gene set (*32, 57, 63*). Gene names in the gene set were converted to their mouse homolog. The Welch Two Sample t-test were used to assess whether the distributions of cell scores between the WT and $\mu$MT cell populations were significantly different. In both cases, the distributions were deemed significantly different with a very low p-value ($p < 2.2 \times 10^{-16}$).

**1000 Immunomes Analyses**
Samples were collected and processed as previously described (*25*). Linear regression was used to regress T cell numbers onto B cell numbers. To determine whether the cell numbers are significantly correlated, p-value of 0.05 from the linear regression in relation to the t-distribution was used.

**NicheNet Analyses:**
Mice: To understand cell-cell interactions associated with age and potential upstream signals driving T cell function by B cells in the aged spleen we used the NicheNetr R package (v.1.0.0) (*23*). Ligand-target prior model, ligand-receptor network, and weighted integrated networks were imported from NicheNet data sets. CD8 T cells clusters (1) and (2), CD4 T cells and CD4 T regs were set separately as receiver/target cell population and both B cell clusters were set as potential senders. The aged spleen was set as the condition at which receiver cells were affected by B cells and the young spleen was set as the reference condition. Then, potential ligands were ranked based on the presence of their target genes in the gene set of interest. The top-20-ranked ligands and top-predicted-target genes of the top-ranked ligands were inferred and visualized in a heatmap and validated on dot plots. All the remaining parameters were default.

Humans: To study the B cell to T cell intercellular communication, we used spleen single-cell datasets from the Tabula Sapiens database (*24*). The .rds file of the "Tabula Sapiens – Immune" dataset was downloaded and imported to Seurat v4. The single-cell RNA-seq data of all three spleen donors were included in the downstream analysis. We performed the analysis using nichenetr R package (v.1.1.0) (*23*) on cells in the dataset belonging to B and T cell types. The expressed genes in sender cells - B cells were selected if they are expressed in at least 5% of the B cell population. The gene set of interest in receiver cells – T cells was defined by adjusted p-value <=0.05 and Log2FC >= 0.25 in the DEGs. Top 32 ligands that were further used to predict activated target genes and construct an activated ligand-receptor network. The other parameters were using default settings.

**Young and Aged B Cell GeneRatio Analyses**
Genes differentially expressed in aged mouse B-cells were obtained from Mogilenko et al. (22) (https://artyomovlab.wustl.edu/sce/var/datasets/Denis/filter/B-cells/files/Aged_vs_Young.tsv). Genes with adjusted P-values < 0.05 and log fold change > 0 were selected for pathway enrichment analysis. These genes were mapped to human orthologs using Ensembl (64) release 103. Pathway enrichment analysis was performed on the human orthologs using pathDIP (65) v.4.0.21.4, with default settings, using all data sources and extended pathways. Enrichment results were visualized for two sets of pathways: antigen presentation pathways and Insulin-PI3K-Akt activity pathways. For each pathway set, enrichment results were visualized as a dot plot with the x-axis indicating GeneRatio (percent of significantly upregulated genes in a pathway), dot color indicating adjusted enrichment P-values (FDR), and dot size indicating the number of significantly upregulated genes in a pathway. Visualization was done in R v.4.0.3, using the ggplot2 package.

**Statistical Analyses:**
Statistical difference between two means was determined via a Mann–Whitney test (i.e., did not assume normal distribution), with GraphPad Prism Software Inc (v.9.0.0) unless otherwise indicated. In figure legends, the n value specified indicates the number of mice, unless stated otherwise. An extreme studentized deviate method (Grubbs' test) was performed to assess for statistical outliers. All data are presented as means ± SEM. Statistical significance was set at $p < 0.05$. * denotes $p < 0.05$, ** denotes $p < 0.01$, and *** denotes $p < 0.001$.

**Table S1 – Primer Sequences**

| Gene | Sequence |
|---|---|
| *TNFA* | Forward: 5′-GTAGCCCACGTCGTAGCAAAC-3′<br>Reverse: 5′-AGTTGGTTGTCTTTGAGATCCATG-3 |
| *INOS* | Forward: 5′-CCGAAGCAAACATCACATTCA-3′<br>Reverse: 5′-GGTCTAAAGGCTCCGGGCT-3′ |
| *IFNG* | Forward: 5’-CGGCACAGTCATTGAAAGCCTA-3’<br>Reverse: 5’- GTTGCTGATGGCCTGATTGTC-3’ |
| *CCL2* | Forward: 5’-CATCCACGTGTTGGCTCA-3’<br>Reverse: 5'-GATCATCTTGCTGGTGAATGAGT-3’ |
| *CCL5* | Forward: 5’-TGCAGAGGACTCTGAGACAGC-3'<br>Reverse: 5’-GAGTGGTGTCCGAGCCATA-3’ |
| *IL1A* | Forward: 5’-TCCATAACCCATGATCTGGAA-3’<br>Reverse: 5’-TTGGTTGAGGGAATCATTCAT-3’ |
| *CDKN2A* | Forward: 5’-AATCTCCGCGAGGAAAGC-3’<br>Reverse: 5’-GTCTGCAGCGGACTCCAT-3’ |
| *CDKN1A* | Forward: 5’-TTGCCAGCAGAATAAAAGGTG-3<br>Reverse: 5’-TTTGCTCCTGTGCGGAAC-3’ |
| *SERPINE1* | Forward: 5’-AGGATCGAGGTAAACGAGAGC-3’<br>Reverse: 5’- GCGGGCTGAGATGACAAA-3’ |
| *INSR* | Forward: 5′- CTGTTCGGAACCTGATGAC -3′<br>Reverse: 5′- ATACCAGAGCATAGGAG -3′ |
| *COL1A1* | Forward: 5′-ACATGTTCAGCTTTGTGGACC-3′<br>Reverse: 5′-TAGGCCATTGTGTATGCAGC-3′ |
| *TGFB1* | Forward: 5′-GGTTCATGTCATGGATGGTGC-3′<br>Reverse: 5′-TGACGTCACTGGAGTTGTACGG-3′ |
| *ACTA2* | Forward: 5′-GGCTCTGGGCTCTGTAAGG-3′<br>Reverse: 5′-CTCTTGCTCTGGGCTTCATC-3’ |
| *ACTB* | Forward: 5’- AGATGTGGATCAGCAAGCAG-3’<br>Reverse: 5’- GCGAAGTTAGGTTTTGTCA-3’ |
| *GAPDH* | Forward: 5’- TCACCACCATGGAGAAGGC-3’<br>Reverse: 5’- GCTAAGCAGTTGGTGGTGCA-3’ |
| *HPRT* | Forward: 5’- CAAGCTTGCTGGTGAAAAGGA-3’<br>Reverse: 5’- TGAAGTACTCATTATAGTCAAGGGCATATC-3’ |

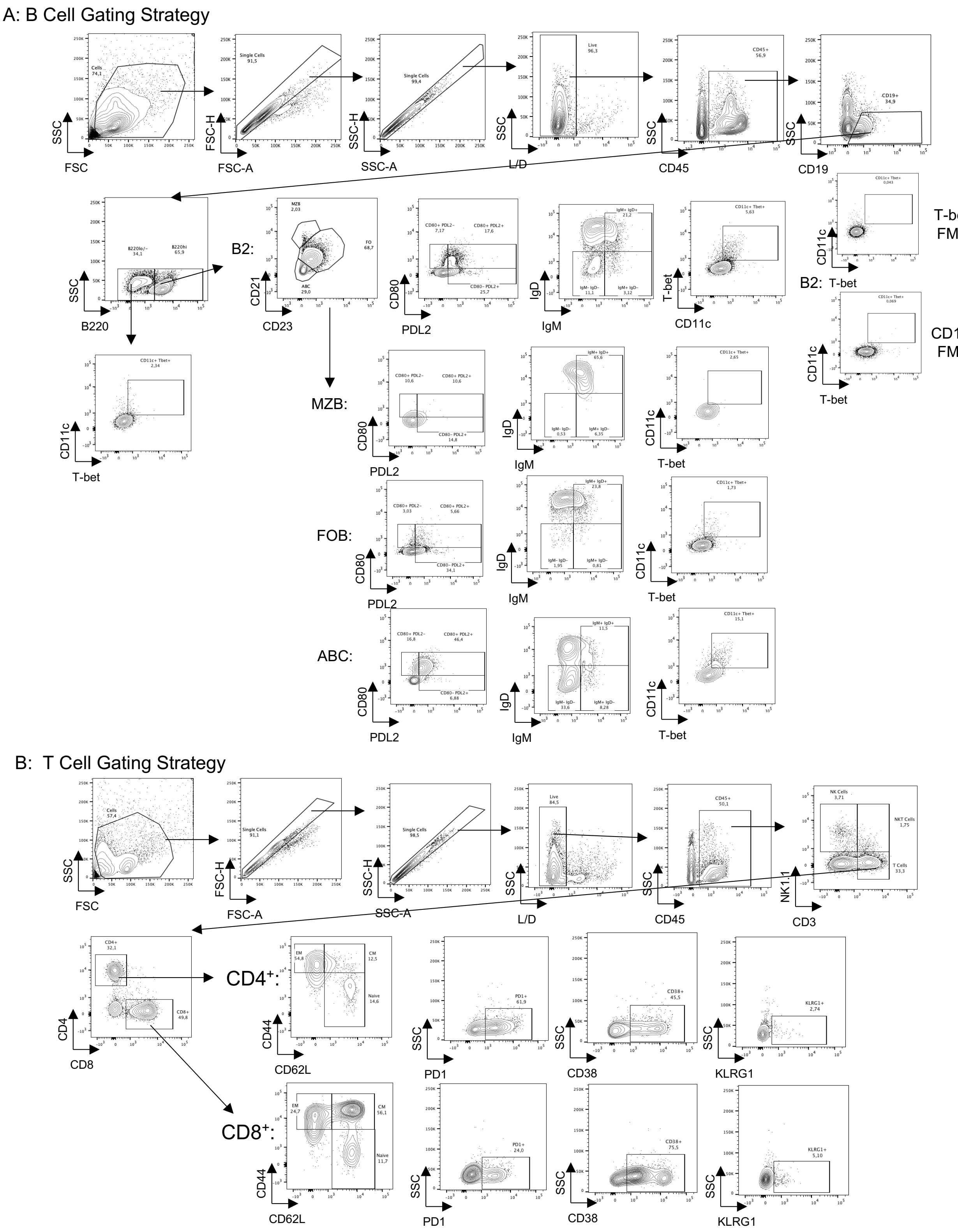


**Fig. S1: Representative gating strategy for assessment of age-related B cells and T cells.** (A) Gating strategy for assessment of age-related B cell compartment. (B) Gating strategy for assessment of age-related T cell compartment.

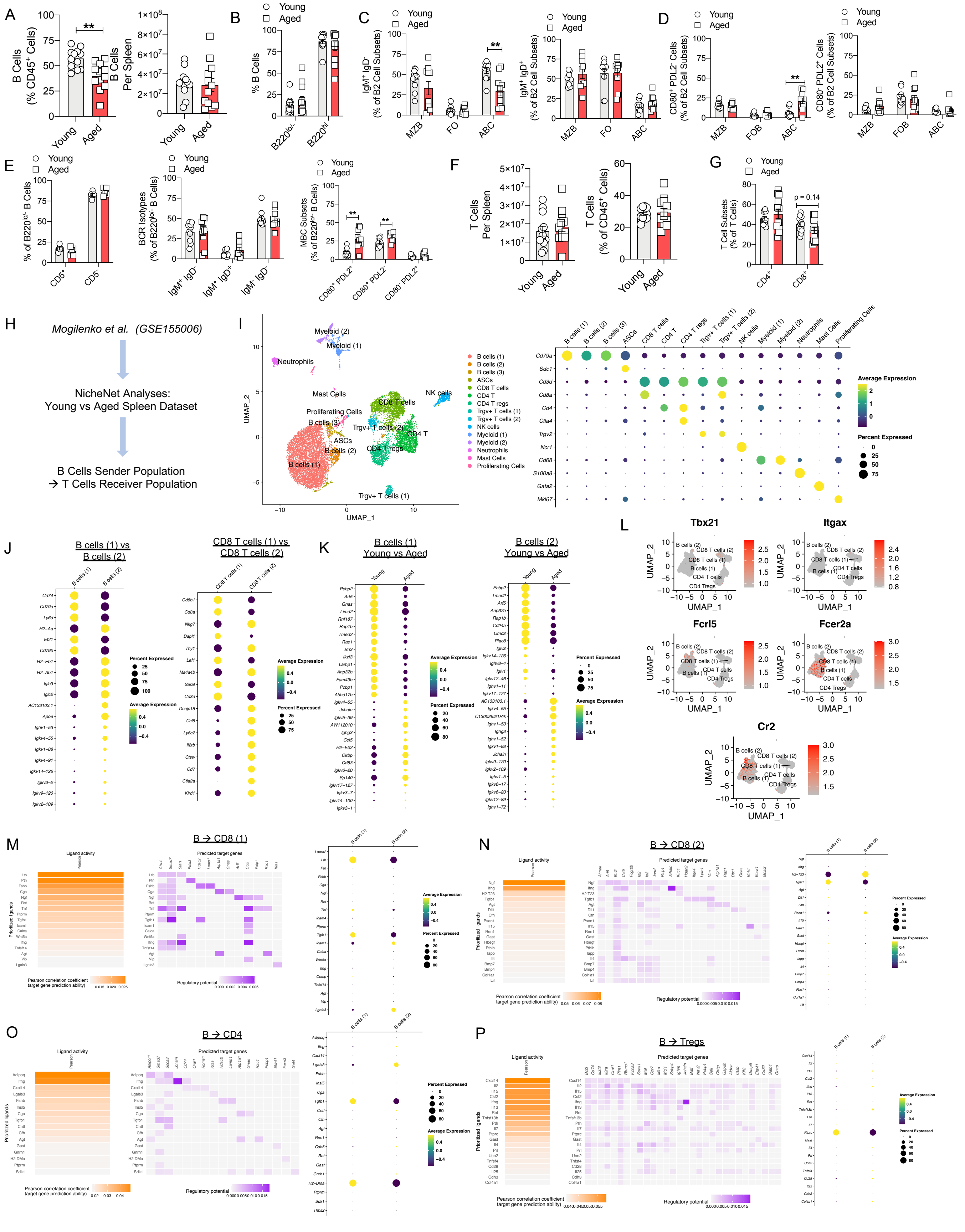


**Fig. S2: Profiling of splenic B and T cells with age.**

(A-G): Supplementary flow cytometric assessment of splenic B cells and T cells in young and aged WT mice (n=5-12 per group). (A) Relative abundance (left) and cell numbers (right) of B cells. (B) Abundance of B cell subsets stratified by B220 expression. (C) Abundance of $IgM^+$ $IgD^-$ (left) and $IgM^+$ $IgD^+$ (right) cells in B2 cell subsets. (D) Abundance of $CD80^+$ $PDL2^-$ (left) and $CD80^-$ $PDL2^+$ (right) cells in B2 cell subsets. (E) Assessment of differential CD5 (left), BCR isotype (middle) and memory B cell (right) subset abundance in B220lo/- B cells. (F) Relative abundance (right) and cell numbers (left) of T cells. (G) Abundance of $CD4^+$ and $CD8^+$ in splenic T cells. (H-P): Supplementary analyses for NicheNet output on young and aged splenic datasets (Mogilenko et al. GSE155006). (H) NicheNet Analysis workflow. (I) UMAP (left) and individual gene expression (right) of all immune cells from aggregated young and aged samples. (J) Differentially expressed genes between re-clustered B cells (1) and (2) (left) and CD8 T cells (1) and (2) right, independent of age-stratification. (K) Differentially expressed genes between young and aged B cells (1) (left) and young and aged B cells (2) (right). (L) Expression UMAP plots of gene markers associated with age-associated B cells. (M) B cell ligand activity (left), T cell gene regulatory potential (middle), and B cell ligand expression levels (right) in the context of B cell clusters influencing cluster CD8 (1). (N) B cell ligand activity (left), T cell gene regulatory potential (middle), and B cell ligand expression levels (right) in the context of B cell clusters influencing cluster CD8 (2). (O) B cell ligand activity (left), T cell gene regulatory potential (middle), and B cell ligand expression levels (right) in the context of B cell clusters influencing cluster CD4. (P) B cell ligand activity (left), T cell gene regulatory potential (middle), and B cell ligand expression levels (right) in the context of B cell clusters influencing cluster Tregs. Data are means ± SEM. *denotes $p < 0.05$, and ** denotes $p < 0.01$.

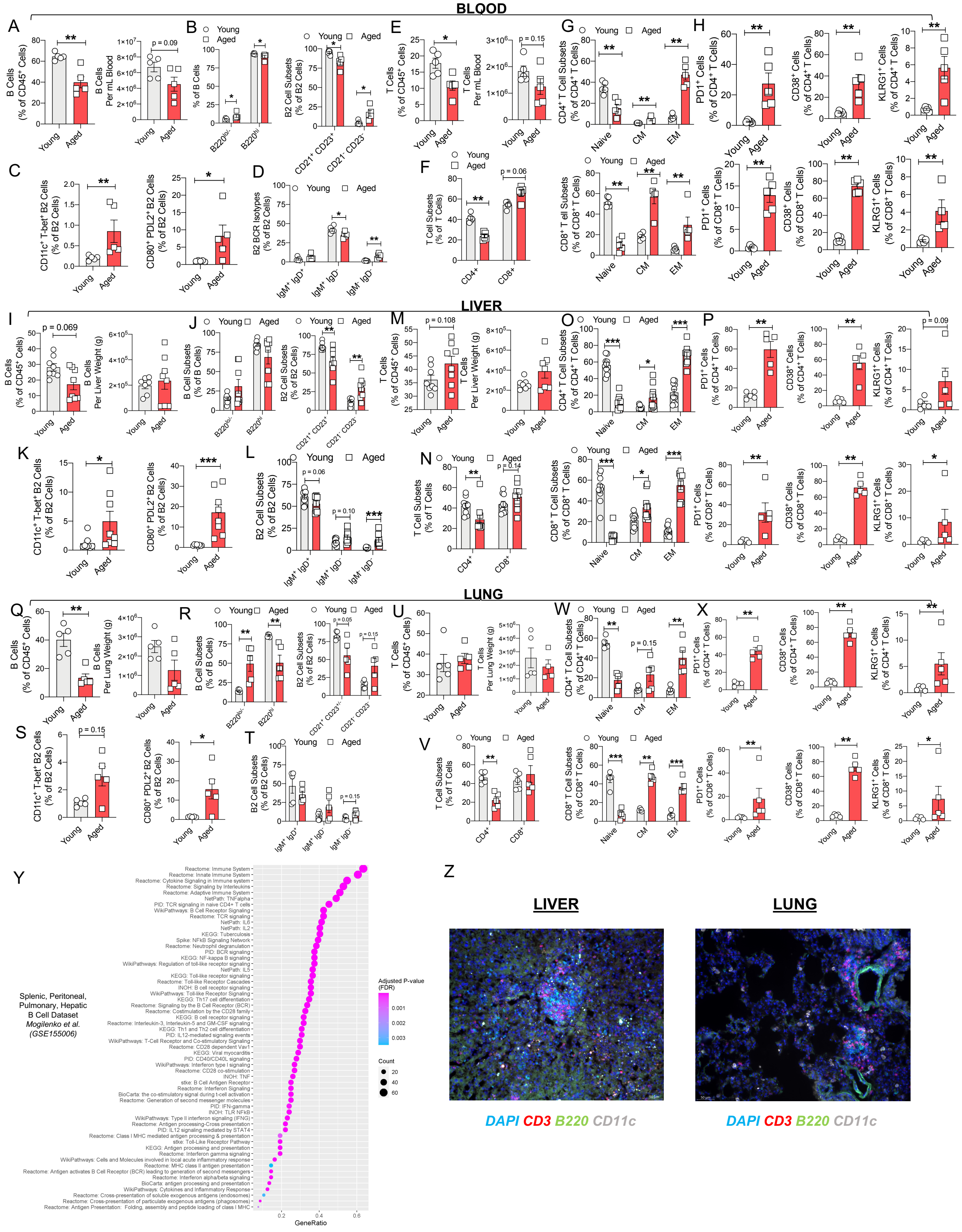


**Fig. S3: Profiling of systemic B and T cells with age.**

(A-H) Flow cytometric assessment B and T cells in blood of young and aged WT mice (n=5 per group). (A) Relative abundance (left) and cell number (right) of total B cells. (B) Abundance of B cell subsets (left) and B2 subsets (right) (C) Abundance of CD11c$^{+}$ T-bet$^{+}$ (left) and CD80$^{+}$ PDL2$^{+}$ (right) cells in B2 cells. (D) Abundance of differential isotypes in B2 cells. (E) Relative abundance (left) and cell number (right) of T cells. (F) Abundance of $CD4^+$ and $CD8^+$ cells in T cells. (G) Abundance of naive, effector memory, and central memory T cells in $CD4^+$ T cells (top) and $CD8^+$ T cells (bottom). (H) Abundance of $PD1^+$ (left), $CD38^+$ (middle), and $KLRG1^+$ (right) cells in $CD4^+$ T cells (top) and $CD8^+$ T cells (bottom). (I-P) Flow cytometric assessment of B and T cells in perfused livers of young and aged WT mice (n=5-8 per group). (I) Relative abundance (left) and cell number (right) of total B cells. (J) Abundance of B cell subsets (left) and B2 subsets (right). (K) Abundance of $CD11c^+$ $T\text{-}bet^+$ (left) and $CD80^+$ $PDL2^+$ (right) cells in B2 cells. (L) Abundance of differential isotypes in B2 cells. (M) Relative abundance (left) and cell number (right) of total T cells. (N) Abundance of $CD4^+$ and $CD8^+$ cells in T cells. (O) Abundance of naive, effector memory, and central memory T cells in $CD4^+$ T cells (top) and $CD8^+$ T cells (bottom). (P) Abundance of $PD1^+$ (left), $CD38^+$ (middle), and $KLRG1^+$ (right) cells in $CD4^+$ T cells (top) and $CD8^+$ T cells (bottom). (Q-X) Flow cytometric assessment of B and T cells in perfused lungs of young and aged WT mice (n=4-5 per group). (Q) Relative abundance (left) and cell number (right) of total B cells. (R) Abundance of B cell subsets (left) and B2 subsets (right). (S) Abundance of $CD11c^+$ $T\text{-}bet^+$ (left) and $CD80^+$ $PDL2^+$ (right) cells in B2 cells. (T) Abundance of differential isotypes in B2 cells. (U) Relative abundance (left) and cell number (right) of total T cells. (V) Abundance of $CD4^+$ and $CD8^+$ cells in T cells. (W) Abundance of naive, effector memory, and central memory T cells in $CD4^+$ T cells (top) and $CD8^+$ T cells (bottom). (X) Abundance of $PD1^+$ (left), $CD38^+$ (middle), and $KLRG1^+$ (right) cells in $CD4^+$ T cells (top) and $CD8^+$ T cells (bottom). (Y) GeneRatio analysis of bulk splenic, hepatic, pulmonary, and peritoneal B cells highlighting activation and antigen presentation related pathways upregulated in aged WT mice compared to young WT mice (Mogilenko et al. GSE155006). (Z) Immunofluorescent staining of CD3, B220 and CD11c in liver (left) and lung (right) from aged WT mice. Data are means ± SEM. * denotes $p < 0.05$, and ** denotes $p < 0.01$.

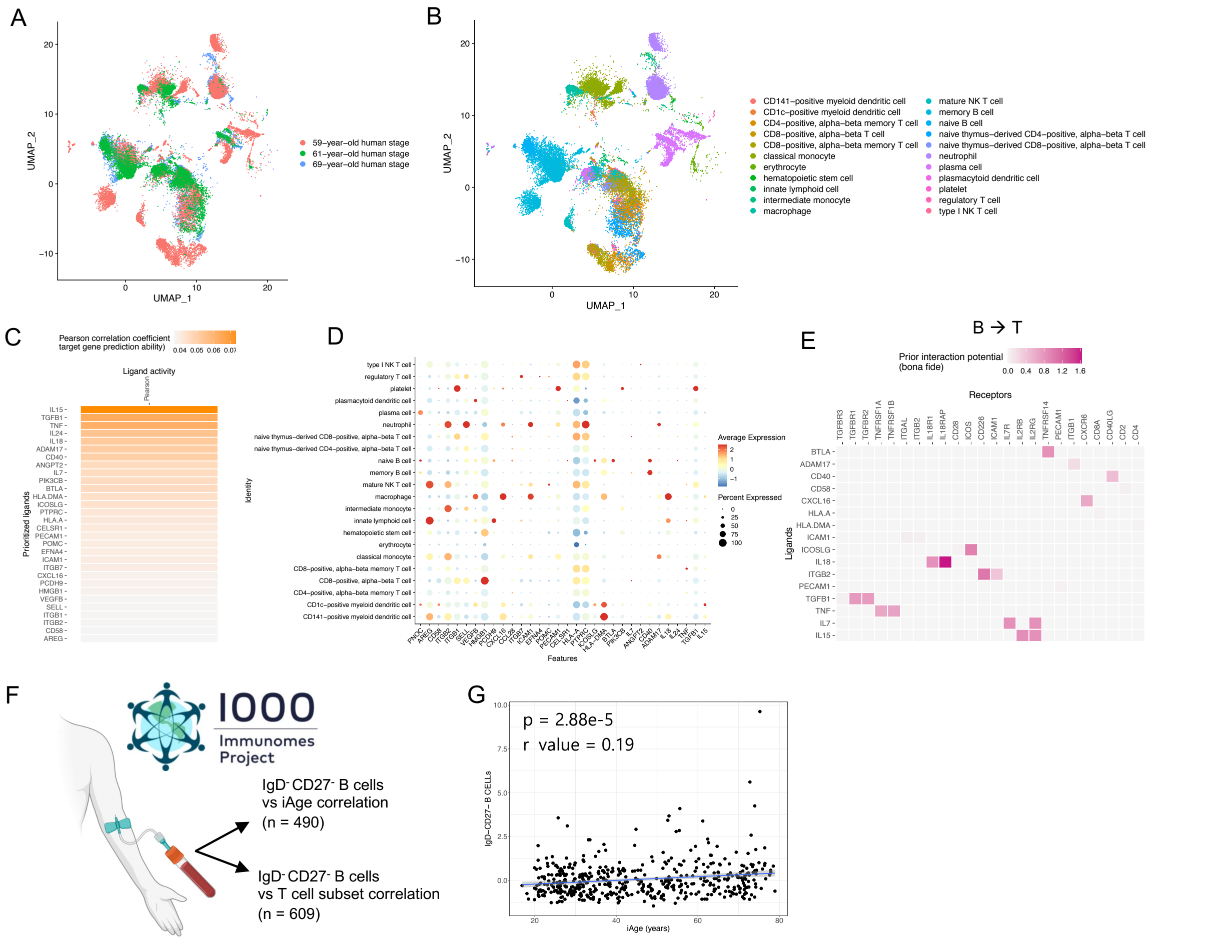


**Fig. S4: Assessment of human B and T cell dysfunction with age.**
(A-E) NicheNet analysis for assessment of B cell and T cell crosstalk in aged human spleens from Tabula Sapiens dataset (The Tabula Sapiens Consortium, GSE201333). (A) UMAP plot highlighting contributions of 3 different donors to cell clusters. (B) UMAP plot with cell annotation from aggregated donor samples. (C) Ligand activity of B cells from aggregated donor sample. (D) Gene expression of key ligands across cell clusters from aggregated donor samples. (E) NicheNet predictive analysis highlighting B cell ligands interacting with T cell receptors during aged conditions, based on cell clusters from aggregated donor samples. (F) Experimental design for assessment of IgD- CD27- ABC-like B cells from 1000 immunomes project. (G) Correlation of iAge with amount of IgD- CD27- B cells in tested subjects (n= 490).

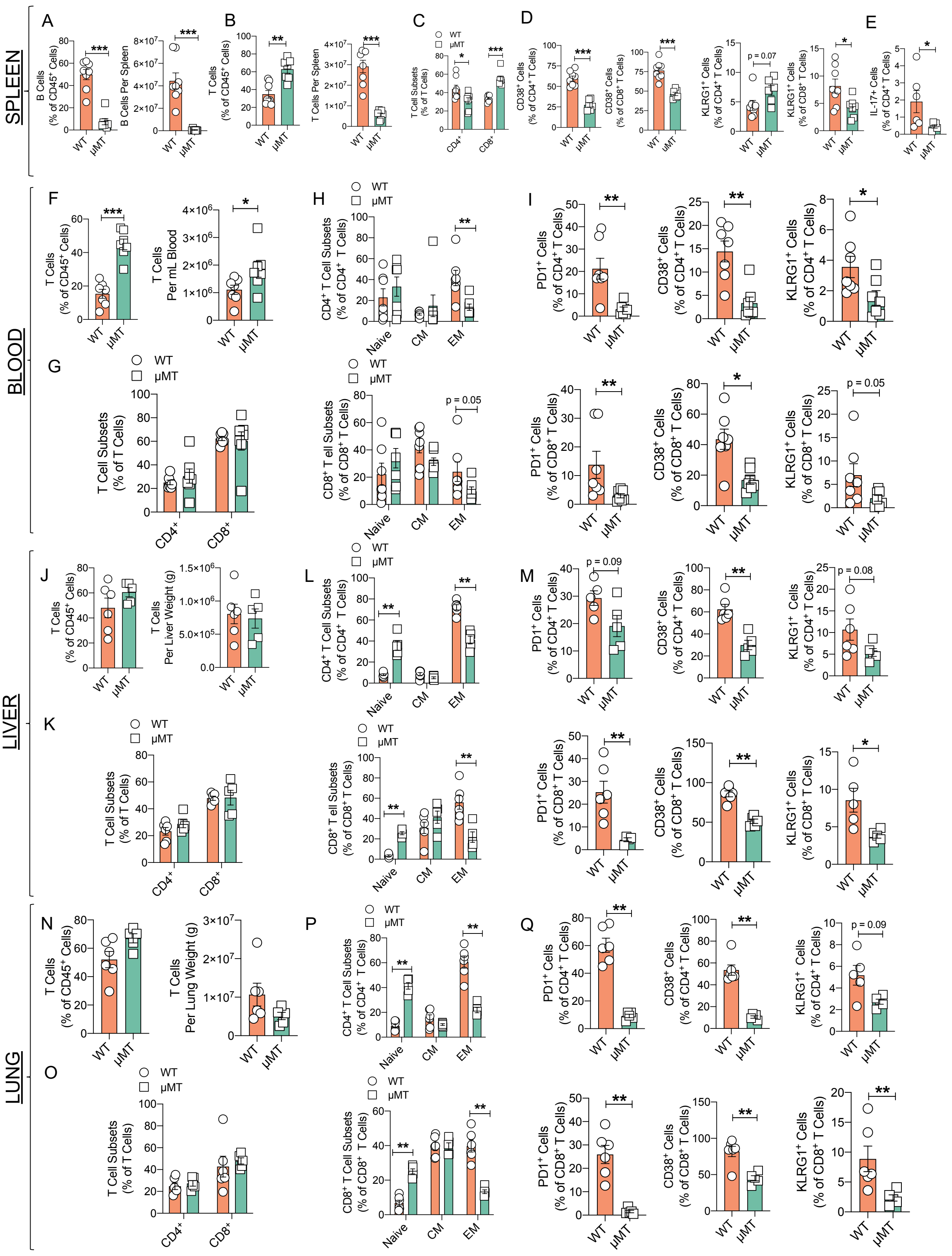


**Fig. S5: Profiling of splenic B cells and systemic T cells in aged μMT mice.**
(A-E) Assessment of splenic B and T cells in aged μMT and WT mice (n=6-8 per group). (A) Relative abundance (left) and cell number (right) of B cells. (B) Relative abundance (left) and cell number (right) of T cells. (C) Abundance of CD4+ and CD8+ cells T cells. (D) Abundance of

$CD38^+$ cells in $CD4^+$ (far left) and $CD8^+$ (middle left) T cells, and $KLRG1^+$ cells in $CD4^+$ (middle right) and $CD8^+$ (far right) T cells. (E) Abundance of IL-$17^+$ cells in $CD4^+$ T cells. (F-I) Assessment of T cells in blood of aged μMT and WT mic (n=7 per group). (F) Relative abundance (left) and cell number (right) of total T cells. (G) Abundance of $CD4^+$ and $CD8^+$ cells in T cells. (H) Abundance of naive, effector memory, and central memory T cells in $CD4^+$ T cells (top) and $CD8^+$ T cells (bottom). (I) Abundance of $PD1^+$ (left), $CD38^+$ (middle), and $KLRG1^+$ (right) cells in $CD4^+$ T cells (top) and $CD8^+$ T cells (bottom). (J-M) Assessment of T cells in perfused livers of aged μMT and WT mice (n=4-6 per group). (J) Relative abundance (left) and cell number (right) of total T cells. (K) Abundance of $CD4^+$ and $CD8^+$ cells in T cells. (L) Abundance of naive, effector memory, and central memory T cells in $CD4^+$ T cells (top) and $CD8^+$ T cells (bottom). (M) Abundance of $PD1^+$ (left), $CD38^+$ (middle), and $KLRG1^+$ (right) cells in $CD4^+$ T cells (top) and $CD8^+$ T cells (bottom). (N-Q) Assessment of T cells in perfused lungs of aged μMT and WT mice (n=5-6 per group). (N) Relative abundance (left) and cell number (right) of total T cells. (O) Abundance of $CD4^+$ and $CD8^+$ cells in T cells. (P) Abundance of naive, effector memory, and central memory T cells in $CD4^+$ T cells (top) and $CD8^+$ T cells (bottom). (Q) Abundance of $PD1^+$ (left), $CD38^+$ (middle), and $KLRG1^+$ (right) cells in $CD4^+$ T cells (top) and $CD8^+$ T cells (bottom). Data are means ± SEM. * denotes $p < 0.05$, ** denotes $p < 0.01$, and *** denotes $p < 0.001$.

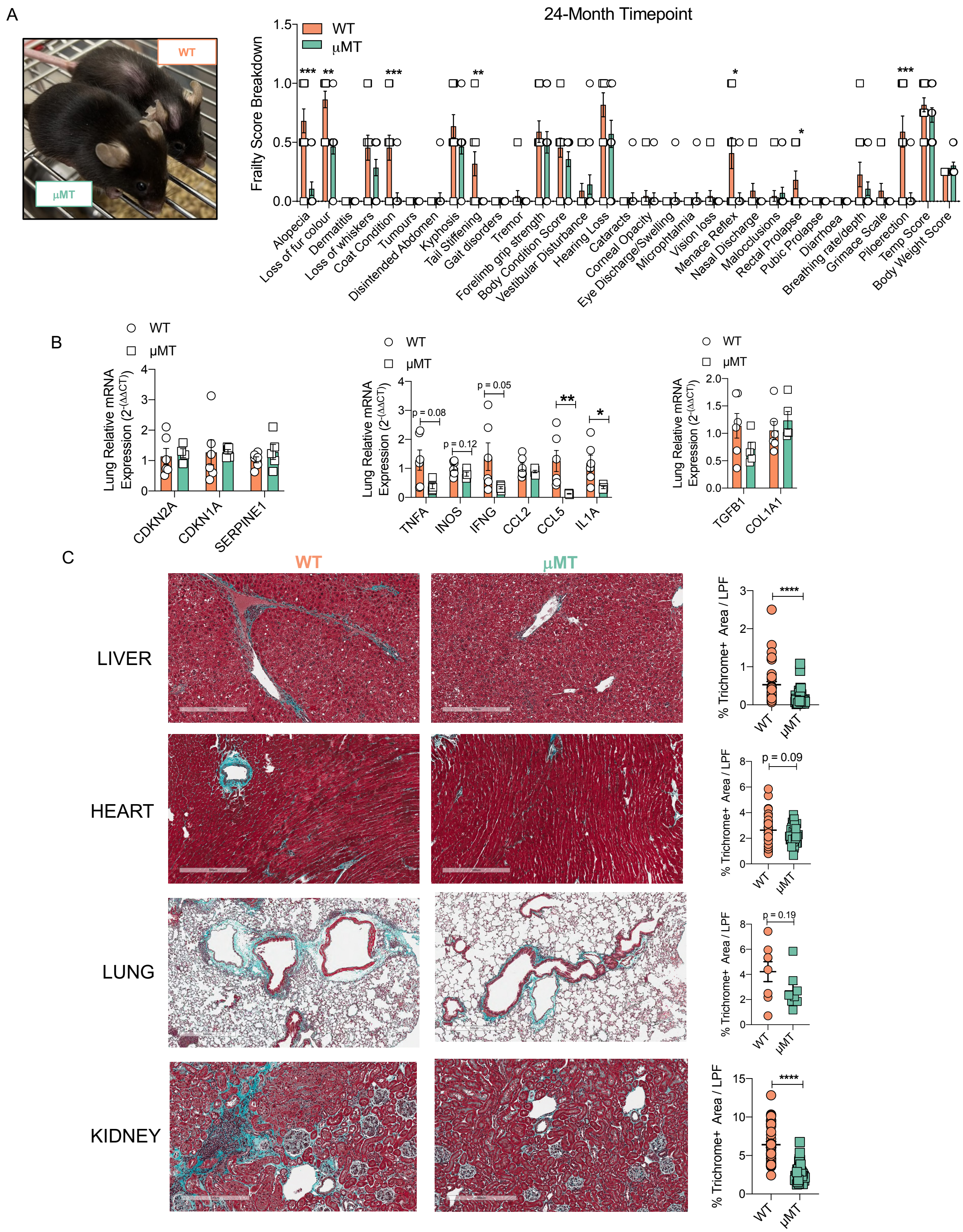


**Fig. S6: Supplemental assessment of healthspan parameters of aged μMT mice.**
(A) Representative image of aged μMT and WT mice (left) and breakdown of 31-index frailty scores in aged μMT and WT mice at the 24-month timepoint (right) (n=11-14 per group). (B) Gene expression of senescence (left), senescence associated secretory phenotype (middle) and fibrosis (right) markers in lungs of aged *μ*MT and WT mice (n=4-6 per group). (C) Representative images (left) and quantification (right) of tissue fibrosis via mason's trichrome

staining (n = 8-40 LPF (tissue dependent) from 4 mice per group). Data are means ± SEM. * denotes $p < 0.05$, ** denotes $p < 0.01$, and *** denotes $p < 0.001$.

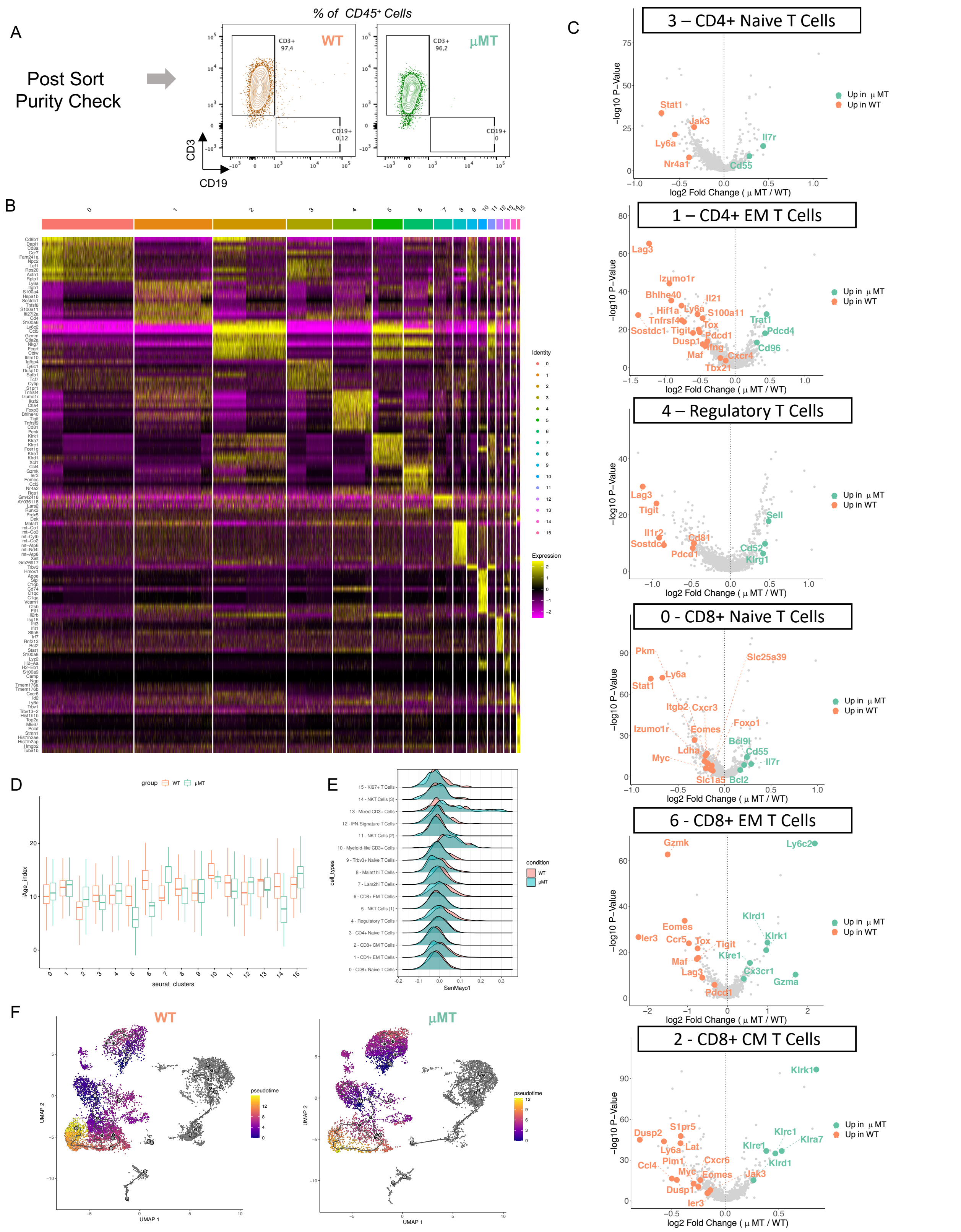


**Fig. S7: Supplemental assessment of T cell transcriptomic changes in aged µMT mice.** (A-F) 5' single cell transcriptomics with VDJ T cell receptor analyses on splenic CD3+ cells from aged uMT and WT mice (pooled n=2 co-housed mice per group). (A) Purity of sorted CD3+ cells as a percentage of total CD45+ cells. (B) Heatmap of top genes per cluster from splenic CD3+ cells from aggregated aged $\mu$MT and WT data. (C) Volcano plot of genes globally upregulated in CD3+ clusters from aged $\mu$MT and WT mice. (D) iAge Index of CD3+ clusters from aged $\mu$MT and WT mice. (E) SenMayo score of $CD3^{+}$ clusters from aged $\mu$MT and WT mice. (F) Pseudotime expression with trajectory analysis of $CD3^{+}$ cells from aged $\mu$MT and WT mice.

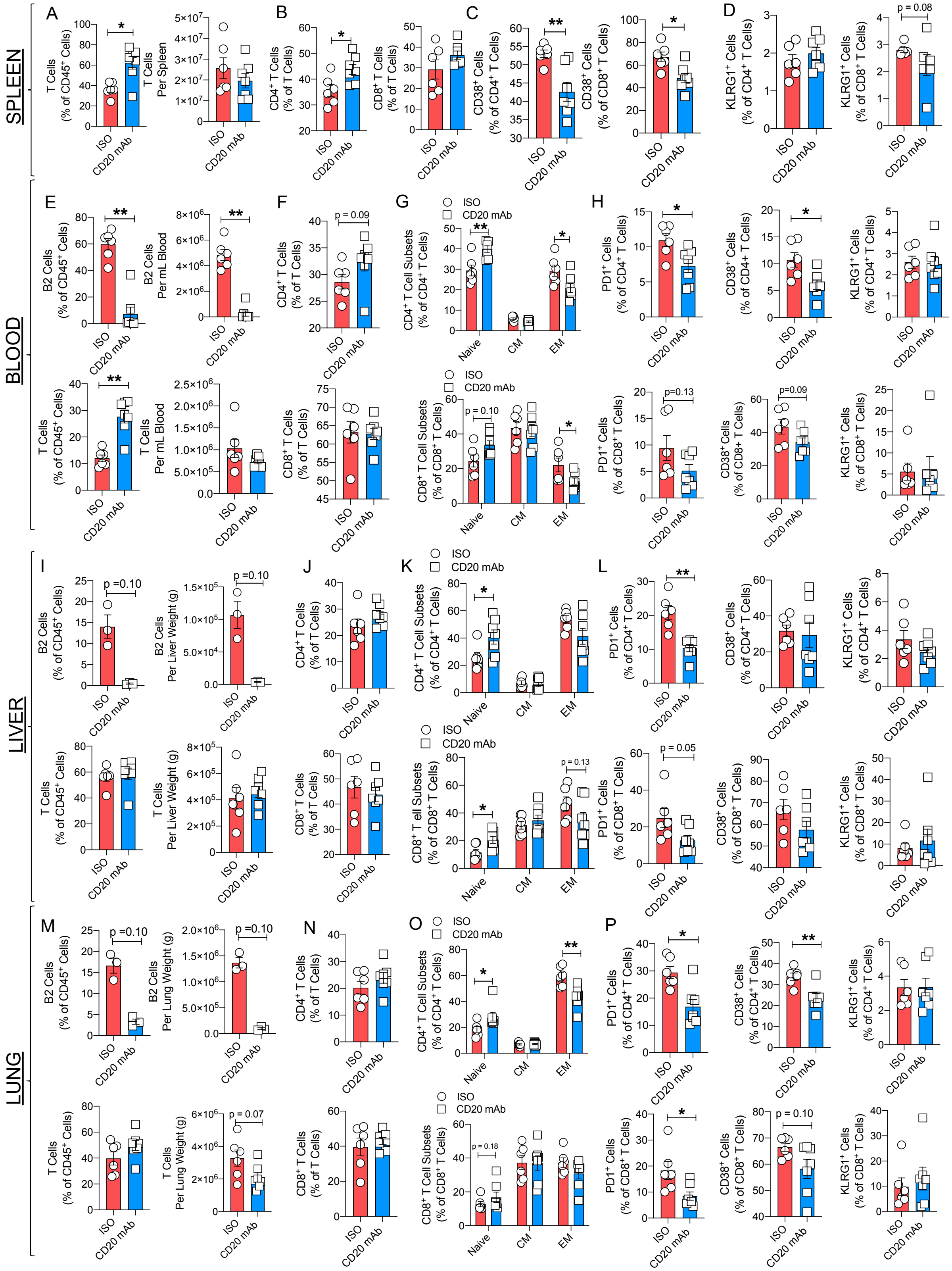


**Fig. S8: Supplemental assessment of the systemic adaptive immune compartment in CD20 mAb treated aging mice.**

(A-D) Assessment of T cells in spleens of aging CD20 mAb and isotype control treated WT mice (n=5-7 per group). (A) Relative abundance (left) and cell number (right) of T cells. (B) Abundance of $CD4^{+}$ (left) and $CD8^{+}$ cells (right) in T cells. (C) Abundance of $CD38^{+}$ cells in $CD4^{+}$ (left) and $CD8^{+}$ (right) T cells. (D) Abundance of $KLRG1^{+}$ cells in $CD4^{+}$ (left) and $CD8^{+}$

(right) T cells. (E-H) Assessment of B2 and T cells in blood of aging CD20 mAb and isotype control treated WT mice (n=6-7 per group). (E) Relative abundance (left) and cell number (right) of B2 cells (top) and T cells (bottom). (F) Abundance of $CD4^+$ (top) and $CD8^+$ cells (bottom) in T cells. (G) Abundance of naive, effector memory, and central memory T cells in $CD4^+$ T cells (top) and $CD8^+$ T cells (bottom). (H) Abundance of $PD1^+$ (left), $CD38^+$ (middle), and $KLRG1^+$ (right) cells in $CD4^+$ T cells (top) and $CD8^+$ T cells (bottom). (I-L) Assessment of B2 and T cells in perfused livers of aging CD20 mAb and isotype control treated WT mice (n=3-7 per group). (I) Relative abundance (left) and cell number (right) of B2 cells (top) and T cells (bottom). (J) Abundance of $CD4^+$ (top) and $CD8^+$ cells (bottom). (K) Abundance of naive, effector memory, and central memory T cells in $CD4^+$ T cells (top) and $CD8^+$ T cells (bottom). (L) Abundance of $PD1^+$ (left), $CD38^+$ (middle), and $KLRG1^+$ (right) cells in $CD4^+$ T cells (top) and $CD8^+$ T cells (bottom) (M-P) Assessment of B2 and T cells in perfused lungs of aging CD20 mAb and isotype control treated WT mice (n=3-7 per group). (M) Relative abundance (left) and cell number (right) of B2 cells (top) and T cells (bottom). (N) Abundance of $CD4^+$ (top) and $CD8^+$ cells (bottom). (O) Abundance of naive, effector memory, and central memory T cells in $CD4^+$ T cells (top) and $CD8^+$ T cells (bottom). (P) Abundance of $PD1^+$ (left), $CD38^+$ (middle), and $KLRG1^+$ (right) cells in $CD4^+$ T cells (top) and $CD8^+$ T cells (bottom). Data are means ± SEM. * denotes $p < 0.05$, and ** denotes $p < 0.01$.

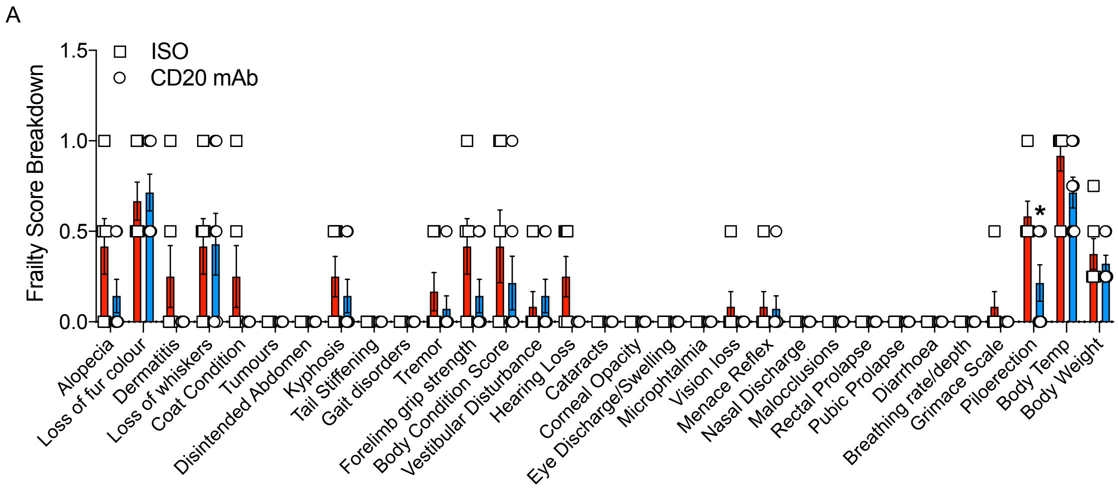


**Fig. S9: Breakdown of 31-index frailty score in CD20 mAb treated aging mice.**
(A) Scoring at the end of treatment (n=6-7 per group). Data are means ± SEM. *denotes $p < 0.05$

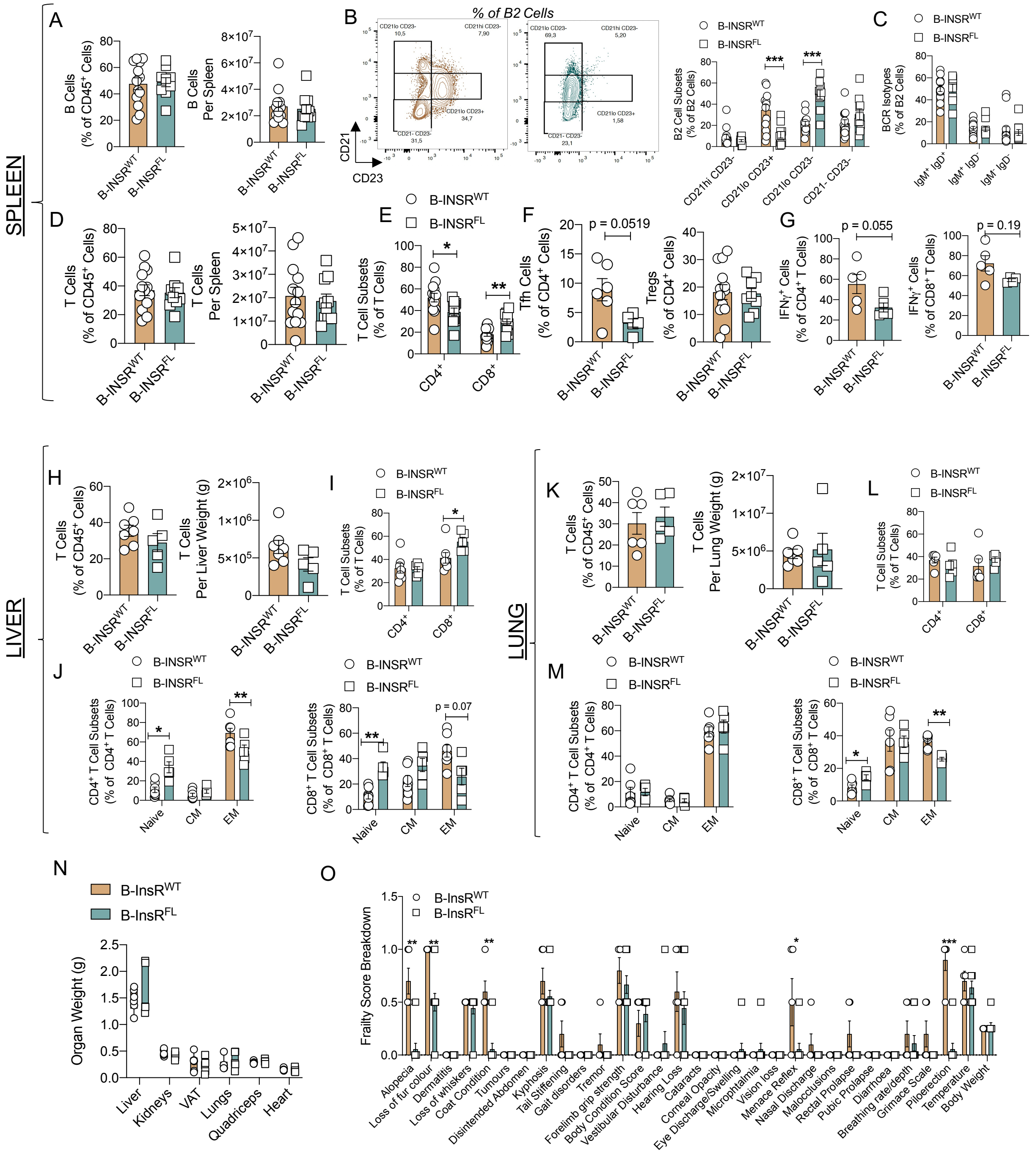

**Fig. S10: Profiling of aged B-InsR$^{FL}$ adaptive immune compartment and healthspan parameters**

(A-G) Assessment of splenic B and T cells in aged B-InsR$^{FL}$ and B-InsR$^{WT}$ mice (n=4-13 per group). (A) Relative abundance (left) and cell numbers (right) of B cells. (B) Representative plots (left) and abundance (right) of B2 cell compartments based on expression of CD21 and CD23. (C) BCR Isotype expression patterns in B2 cells. (D) Relative abundance (left) and cell numbers (right) of T cells. (E) Abundance of $CD4^+$ and $CD8^+$ cells in T cells. (F) Abundance of T follicular helper cells (left) and regulatory T cells (right). (G) Abundance of $IFN\gamma^+$ cells in

$CD4^+$ (left) and $CD8^+$ (right) T cell subsets. (H-J): Assessment of T cells from perfused livers of aged B-InsR$^{FL}$ and B-InsR$^{WT}$ mice (n=5-7 per group). (H) Relative abundance (left) and cell numbers of T cells (right). (I) Abundance of $CD4^+$ and $CD8^+$ cells in T cells. (J) Abundance of naive, effector memory, and central memory T cells in $CD4^+$ T cells (left) and $CD8^+$ T cells (right). (K-M): Assessment of T cells from perfused lung of aged B-InsR$^{FL}$ and B-InsR$^{WT}$ mice (n=4-6 per group). (K) Relative abundance (left) and cell numbers (right) of T cells. (L) Abundance of $CD4^+$ and $CD8^+$ cells in T cells. (M) Abundance of naive, effector memory, and central memory T cells in $CD4^+$ T cells (left) and $CD8^+$ T cells (right). (N) Organ weight of aged B-InsR$^{FL}$ and B-InsR$^{WT}$ mice (n=5-6 per group). (O) 31-index frailty score breakdown of aged B-InsR$^{FL}$ and B-InsR$^{WT}$ mice at the 24-month timepoint (n=5-9 per group). Data are means ± SEM. * denotes $p < 0.05$, ** denotes $p < 0.01$, and *** denotes $p < 0.001$.
